# Regulatory T cells restrain cytotoxic and bystander CD8 T cells without compromising antigen-driven memory in mucosal tissue

**DOI:** 10.64898/2026.02.25.708034

**Authors:** Irene Cruz Talavera, Jessica B. Graham, Jessica L. Swarts, Brianna R. Traxinger, M. Quinn Peters, Lakshmi Warrier, Amanda L. Koehne, Tanvi Arkatkar, Keith R. Jerome, Martin Prlic, Jennifer M. Lund

**Affiliations:** Vaccine and Infectious Disease Division, Fred Hutchinson Cancer Center, Seattle, WA, USA; Department of Global Health, University of Washington, Seattle, WA, USA; Experimental Histopathology, Fred Hutchinson Cancer Center, Seattle, WA, USA; Department of Laboratory Medicine and Pathology, University of Washington, Seattle, WA, USA

## Abstract

Many pathogenic human infections enter the host via a mucosal surface. These nonlymphoid tissues are abundantly populated by polyclonal memory CD8 T cells that persist following infections for protection upon repeat exposure. Memory T cells can be triggered via T cell receptor recognition of their cognate antigen upon re-infection to exert effector functions, including cytotoxicity and cytokine production, and assist in pathogen elimination.

Alternatively, some T cells are ‘bystander activated’ by cytokines without an antigenic signal. This layered approach boosts efficient pathogen clearance but also poses a threat to host tissues if this response is not properly controlled. Here, we investigate the regulatory mechanisms modulating the tissue memory CD8 T cell response upon recall, leveraging viral rechallenge mouse models to distinguish antigen-driven versus cytokine-activated memory tissue CD8 T cell immunity. We and that regulatory T cells (Treg) participate in restricting cytotoxic and bystander activity without compromising the antigen-driven protective memory CD8 T cell response in mucosal T cells. Critically, Treg provide extrinsic regulation of tissue CD8 T cell cytotoxicity in part through restriction of available IL-2 and IL-15 trans-presentation. Our findings help define the extrinsic environmental and cellular cues in mucosal tissues that direct tissue-memory CD8 T cells.

## INTRODUCTION

The tissue microenvironment plays an influential role in guiding phenotypic and functional aspects of lymphocytes such that mucosal tissue T cells have distinct qualities compared to their circulating counterparts^1–3^. As a notable example, tissue-resident memory T cells (Trm) have been identified as a subset of memory T cells with a unique transcriptional signature that persists, patrols, and self-renews within non-lymphoid tissues^3–7^. Upon recognition of their cognate antigen via their TCR, Trm can accelerate protection *in situ* by directly killing infected cells via cytolytic function and through secretion of pro-inflammatory cytokines such as IFNγ, TNFα, and IL-2^1,5,8,9^. This cytokine production not only elicits a rapid local anti-viral response but also serves as a ‘sense and alarm’ function^5^ that potentiates robust tissue immunity through recruitment and activation of other immune cells^8,10,11^. Thus, in addition to canonical antigen-driven memory CD8 T cell activity elicited by pathogen re-exposure in the tissue, memory CD8 T cells can also become bystander-activated via cytokine signals to acquire cytotoxic functions^12–19^. It was recently shown that at steady state, human memory CD8 T cell cytotoxicity, including expression of granzyme B, is lower in nonlymphoid tissues compared to the circulation, and inversely correlates with CD8 Trm phenotype^20^. Nevertheless, exposure of human tissue memory CD8 T cells to IL-15 is sufficient to increase expression of Granzyme B and perforin, each of which is tightly correlated with target cell killing capacity^20^. Thus, while human tissue memory CD8 T cells maintain a state of relative quiescence at steady-state, signals within the tissue environment, such as those elicited during infection, lead to rapid acquisition of cytotoxic function.

However, a knowledge gap remains in terms of how the cytotoxic activity of tissue memory CD8 T cells is controlled, particularly in settings of active inflammation or infection. Moreover, the overall Trm response is clearly potent and effective at controlling pathogens upon re-exposure, yet the mechanisms involved in regulating this potent tissue memory response remain unclear. We predicted that the regulation of this robust and rapid tissue memory T cell response is critical to avoid excessive collateral damage to host tissues. Furthermore, we hypothesized that regulatory T cells (Treg) could provide a layer of cell-extrinsic regulation of tissue T cells in settings of mucosal re-infection.

Treg are a CD4+ T cell subset characterized by their expression of the forkhead box protein 3 (FoxP3) transcription factor and known for their role in regulating the effector function and activation of other immune cells via various direct and indirect mechanisms^21–26^. Though Treg are crucial in promoting tolerance and preventing autoimmunity, whether they help or hinder protective immunity and disease outcomes during infection seems to be dependent on the tissue microenvironment and invading pathogen^27–31^. Treg have been shown to limit immune-mediated pathology by restraining the magnitude and intensity of adaptive immune responses, but often at the expense of timely pathogen clearance^27,29,32^. Early studies utilizing mouse infection models of *M. tuberculosis*, *L. major*, HSV-1, Friend virus, LCMV, and *Plasmodium* demonstrated that inhibition or depletion of immunosuppressive Treg function results in robust protective antigen-specific T cell responses^27–29,32–40^. However, in the context of primary infection within barrier tissues, such as the lung and vaginal tract, Treg are necessary for orchestrating the development of an appropriate anti-viral response^31,41–43^. During respiratory syncytial virus (RSV) infection in mice, *in vivo* depletion of Treg prior to primary infection resulted in hampered recruitment of RSV-specific CD8 T cells to the lung early in infection with consequent exacerbation in disease, morbidity, and delayed viral clearance^42^. Similarly, in a model of intravaginal viral infection in mice, systemic depletion of activated CTLA-4+ Treg during primary herpes simplex virus type 2 (HSV-2) infection in mice caused early death and increased viral titers due to delayed arrival of innate immune cells, impaired priming and early migration of HSV-2-specific CD4+ T cells to the site of infection, and decreased antiviral cytokine production in the vaginal tract^43^. In primary viral infection studies, FoxP3+ Treg rapidly proliferate and accumulate in draining secondary lymphoid organs (SLOs) and traffic to mucosal infection sites (i.e. lung or vaginal tract) with kinetics similar to that of CD4+ conventional T cells (Tconv)^31,42,43^. We previously demonstrated that regulatory T cells (Treg) accumulate within infected tissue and coordinate early immune responses and T cell priming during primary infection^31,43^, though their role in shaping tissue memory T cell responses remains unclear. Furthermore, these mucosal Treg are highly activated based on increased expression of activation and immunosuppressive markers^44,45^. Notably, this influx of activated Treg has also been observed in human studies focused on tissue T cell responses within biopsies of HSV skin lesions. Higher Treg:CD4+ Tconv and Treg:CD8+ T cell ratios within HSV-2 lesions correlated with higher viral loads, and Treg persisted alongside HSV-2-specific Trm throughout lesion healing^46^. However, whether the presence of these Treg are the cause of increased viral replication or a response to robust tissue immunity resulting from viral reactivation is unclear^44,46^. In mice, Treg have been implicated in the suppression of HSV-1 vaccine-generated memory CD8+ T cell responses^34^, leading us to speculate that they may play a similar role in restraining the activation and function of tissue recall responses.

We sought to interrogate the role of Treg in the regulation of the tissue memory T cell response to re-infection. Here, using a mouse model of genital HSV-2 infection, we demonstrate that transient Treg depletion during a tissue recall response to secondary infection contributes to exacerbating tissue pathology following vaginal HSV-2 challenge, despite normal viral clearance. Furthermore, we demonstrate that Treg selectively restrict the CD8 T cell cytotoxic program and bystander activation while sparing the protective memory HSV-specific tissue CD8 T cells in mucosal tissue. This selective Treg-mediated restriction of the cytotoxic activity of tissue memory CD8 T cells is dependent, at least in part, upon both IL-2 availability in the tissue microenvironment and IL-15 trans-presentation by tissue innate myeloid immune cells. Our findings highlight the essential role of Treg in regulating the CD8 T cell memory recall response and bystander-activated cytotoxic lymphocytes within barrier tissue sites, preserving an appropriate and protective pathogen-specific tissue memory T cell response while limiting cell-mediated immunopathology.

## RESULTS

### Primary vaginal HSV-2 infection induces a highly activated Treg response that persists within the vaginal tract

We have previously shown that human and murine vaginal Treg are highly activated at steady state and acquire a phenotypic and transcriptional tissue signature distinct from baseline following primary infection with HSV-2^45^. To address whether Treg in the mucosal tissue could be positioned to play a role in the memory response upon re-exposure to HSV-2, we infected mice intravaginally with thymidine kinase-deficient (TK-) HSV-2 and allowed them to recover for approximately one month. Unlike WT HSV-2, mice survive and recover from intravaginal TK-HSV-2, allowing us to assess the immune response after acute mucosal viral infection^47,48^. To characterize and define transcriptional and phenotypic changes in vaginal Treg at a memory timepoint following primary HSV-2 infection, Treg (CD4+ FoxP3^eGFP^+) were FACS-sorted from the vaginal tract (VT) and draining lymph nodes (dLN; inguinal and iliac) of naïve and previously HSV-2-infected mice for bulk RNA sequencing (Fig. 1A). Compared to Treg isolated from naïve mice, vaginal tissue Treg from HSV-2-infected mice displayed elevated transcripts for canonical markers of immunosuppressive function (*Il10, Ctla4, Icos*), activation (*Cd69, Tnfrsr4*), and tissue-homing receptors (*Ccr8*). Although the subset of differentially expressed genes between vaginal tissue Treg from naïve and previously infected mice was relatively small, this data suggests that infection-experienced VT Treg retain heightened activation, tissue homing ability, and suppressive potential long after resolution of acute infection. Furthermore, there were markedly fewer differentially expressed genes between Treg from dLN of naïve and HSV-2-infected mice. We next assessed Treg abundance and phenotypes at day 7 and day 90 after HSV-2 infection using flow cytometry (Fig. S1). As previously demonstrated, Treg numbers increased significantly both within the vaginal tissue and dLN at day 7 post-infection (p.i.)^31,45^. However, at day 90 p.i. Treg numbers remained elevated compared to baseline in the vagina but not the dLN (Fig. 1B). In addition to Treg abundance, we assessed markers of activation and immunosuppressive function. Vaginal tissue Treg displayed significantly increased frequencies of CD25, CTLA-4, ICOS, Tim-3, and GITR out to day 90 post-HSV-2 infection, whereas dLN Treg only transiently expressed these markers during acute infection (Fig. 1C), consistent with transcriptional profiling that showed more persistent changes in VT Treg compared to dLN Treg after infection (Fig. 1A). Thus, highly activated Treg accumulate within the vaginal tissue, remain transcriptionally and phenotypically activated, and persist long after clearance of mucosal viral infection, potentially poised to participate in governance of the recall tissue T cell response.

**Figure 1.**
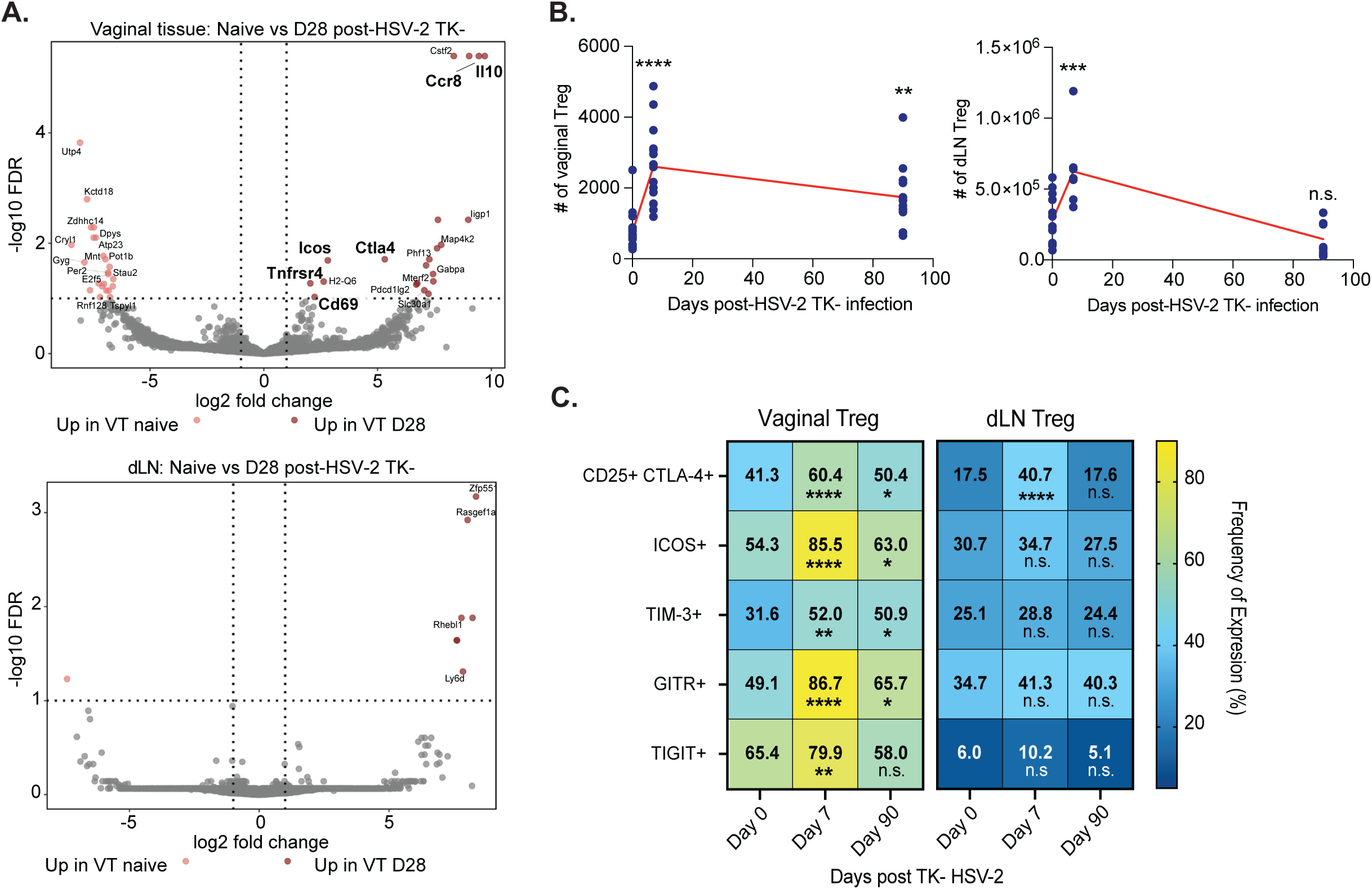
Primary vaginal HSV-2 infection induces a highly activated Treg phenotype that persists within the vaginal tract. Depo Provera-treated FoxP3^eGFP^ mice were intravaginally infected with attenuated thymidine kinase-deficient HSV-2 (TK-) and harvested for dLN and vaginal tract (VT) tissues on day 28 post-infection (n=4). Naïve Depo-Provera-treated mice were included as controls (n=4). Samples were prepared for fluorescence-activated cell sorting (FACS). CD4+ FoxP3+ Treg were sorted from dLN and VT single-cell suspensions and processed for bulk RNA-sequencing. For comparison of differentially expressed genes, genes with a False Discovery Rate (FDR) of less than 0.1 and an absolute expression fold-change of greater than 1.0 were considered differentially expressed. (**A**) Volcano plots show significantly differentially expressed genes between tissues from naïve vs. infected mice (VT top, dLN bottom). (**B**) VT and dLN were collected at 7 (acute timepoint) and 90 (memory timepoint) days post-infection and processed for flow cytometric analysis, with naïve Depo Provera-treated mice included as day 0 uninfected controls. Absolute numbers of CD4+ FoxP3+ Treg in the VT (left) and dLN (right) at day 0, 7, and 90 are shown. (**C**) Frequencies of Treg expressing markers of activation and immunosuppressive function in the VT (left) and dLN (right) at day 0, 7, and 90. Data shown in B and C are combined from two experiments per time point (n= 3-7 mice per group). Error bars represent the mean and SD. Statistical significance was determined by One-Way ANOVA and Dunnett’s multiple comparisons test.

### Treg limit immune-mediated tissue pathology upon HSV-2 re-infection

Having confirmed the persistence of Treg in the vagina at a memory time point after the resolution of primary HSV-2 infection, we wanted to assess the functional impact of Treg on disease progression and pathogen clearance upon HSV-2 challenge (Fig. 2A)^49^. FoxP3^DTR^ mice were first infected with TK-HSV-2 and allowed to recover. One month after primary infection, mice were systemically depleted of Treg, resulting in depletion of dLN and vaginal Treg (Fig. 2B), and subsequently challenged with WT HSV-2. Notably, the frequency and number of FoxP3+ Treg in the VT was significantly higher in FoxP3^WT^ mice 3 days post-challenge than in unchallenged mice (about a month after primary TK-HSV-2), demonstrating the expansion or recruitment of Treg upon re-infection (Fig. 2B; S2A). To assess the impact of Treg depletion on disease progression upon challenge, vaginal tissue was harvested from Treg-sufficient and depleted mice 7 days post-challenge (p.c.), formalin-fixed, paraffin-embedded, H&E-stained, and blindly scored by a pathologist for tissue damage, signs of inflammation, and lymphocyte infiltration. We utilized a total sum histopathology scoring system in which different regions of the VT (lumen, epithelium, lamina propria, and muscularis) were individually scored from 0 to 4, summed, and averaged for a total score per mouse^50^ (Fig. 2C). Daily vaginal washes were also collected from challenged mice up to the time of harvest to assess changes in viral load via RT-PCR^19^. Naïve Treg-sufficient and depleted mice were included as controls for inflammation caused by the transient systemic ablation of Treg^51^.

**Figure 2.**
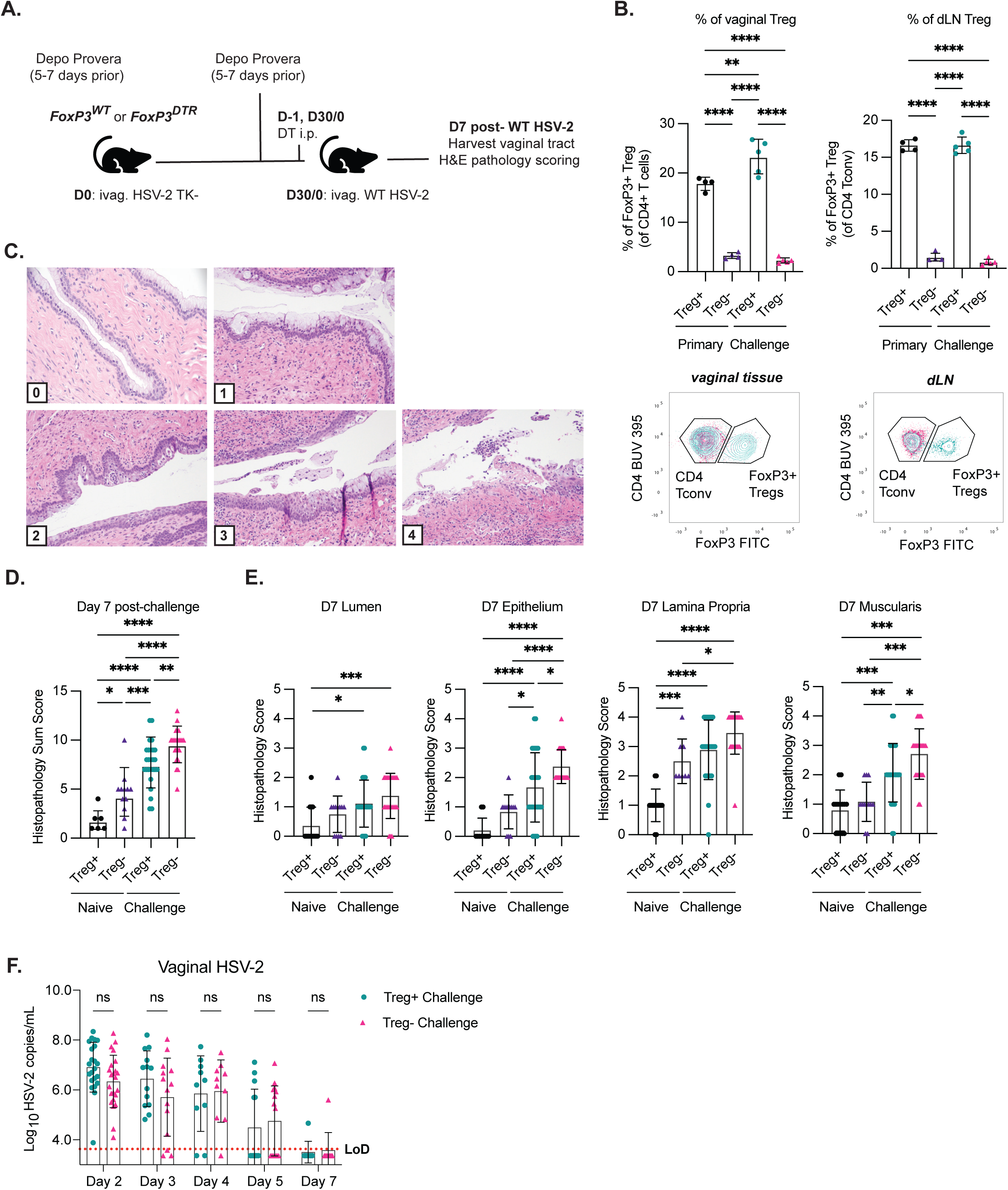
Treg limit immune-mediated tissue pathology upon HSV-2 re-infection. (**A**) Female FoxP3^WT^ and FoxP3^DTR^ Depo Provera-treated mice were vaginally infected with 10^5^ PFU of attenuated HSV-2 TK-and allowed to recover for 28-30 days. Mice were again treated with Depo Provera 5-7 days before WT HSV-2 (10^4^ PFU) challenge and treated intraperitoneally with diphtheria toxin (DT) on days-1 and 0 (with respect to challenge) to ablate Treg prior to infection (FoxP3^DTR^ mice, Treg-depleted group, Treg-). A dose of 30 μg/kg was administered the day before WT challenge, followed by a lower dose of 10 μg/kg on d0. (**B**) VT and dLN were harvested at 3 days p.c. from unchallenged (primary TK-infection only) and WT HSV-2-challenged mice and characterized via flow cytometry to assess Treg depletion (Treg+ challenged group in green, Treg-challenged group in pink). Representative data from one of two experiments (n= 4-5 mice per group) are shown. VT (left) and dLN (right) frequencies of FoxP3+ Treg in Treg-depleted and sufficient mice after WT HSV-2 challenge and representative staining of FoxP3+ CD4+ Treg in Treg-depleted and sufficient mice after WT HSV-2 challenge. Statistical significance was determined by One-Way ANOVA and Dunn-Šídák multiple comparisons tests. (**C**) Female FoxP3^WT^ and FoxP3^DTR^ mice were infected and treated with DT. VT tissues were carefully harvested at a week p.c., formalin-fixed, paraffin-embedded, and scored blindly by a pathologist using a total sum pathology rubric. An example of pathology scoring (0-4) in H&E-stained vaginal epithelium. Naïve Treg-sufficient and Treg-depleted mice were included as controls for tissue damage caused by DT Treg depletion alone. (**D**) Total sum vaginal pathology scores on day 7 p.c. (**E**) Total sum pathology is composed of averaged and summed scores within the lumen, mucosal epithelium, lamina propria, and muscularis mucosa layers of the VT. Data combined from 2-3 representative experiments (n = 3-7 mice per group). Error bars represent the mean and SD. Statistical significance was determined by Two-Way ANOVA and Tukey’s multiple comparisons test. (**F**) Vaginal washes were collected daily during the first week of infection and assessed for viral load via RT-PCR. Data shown are combined from 3 representative experiments per time point. Error bars represent the mean and SD. Statistical significance was determined by multiple unpaired T-tests with Welch correction and Holm-Šídák multiple hypothesis testing.

Mice challenged with WT HSV-2 did not show external signs of infection or disease progression as assessed by clinical scoring^49^, irrespective of Treg depletion. Treg depletion alone in naïve mice resulted in a significant increase in vaginal sum pathology scores, though to a lesser extent compared to the scores quantified 7 days after HSV-2 challenge (Fig. 2D). Further, Treg-depleted and challenged mice displayed significantly higher total vaginal sum pathology scores compared to Treg-sufficient and challenged mice at 7 days p.c., with significantly higher average scores within the vaginal epithelium and muscularis (Fig. 2D-E). Despite this increase in pathology scores, we did not observe a difference in vaginal viral load or ability to clear the infection (Fig. 2F), suggesting that the tissue damage was likely immune-mediated and not due to an increase in viral burden in the absence of Treg as observed in primary infection^31^. Critically, this indicates that Treg functionally spare the protective tissue immune response and demonstrates that Treg are essential to manage a protective tissue recall response that limits host tissue pathology.

### Treg limit the magnitude and cytotoxic potential of the CD8 T cell recall response shortly after WT HSV-2 challenge

We next sought to assess how Treg impact the recall memory CD8 T cell response shortly after WT HSV-2 challenge (Fig. 3A). Groups of Foxp3^WT^ or Foxp3^DTR^ mice received a primary TK-HSV-2 infection, and one month later, were treated with DT and challenged with WT HSV-2 followed by collection of vaginal and dLN tissues at 3 days (p.c.); or included as controls, unchallenged Foxp3^WT^ or Foxp3^DTR^ mice following primary infection only, but at a memory timepoint matched to mice that received vaginal HSV-2 challenge. Unchallenged primary HSV-only groups were also treated with Depo-Provera at the same time points to control for differences in the estrus cycle between groups. We first evaluated the abundance and phenotypic changes of CD8+ CD3e+ T cells in dLN and VT shortly after challenge using flow cytometry. Treg-depleted and challenged mice had a significantly higher number of CD8 T cells in the vaginal tract at day 3 post-challenge (p.c.) compared to Treg-sufficient challenged mice; this significant increase in number was not observed in dLN CD8 T cells (Fig. 3B and Fig. S2B). In alignment with their increased number, a statistically larger frequency and number of vaginal and dLN CD8 T cells from Treg-depleted and challenged compared to Treg-sufficient and challenged mice expressed Ki67 and CD44, indicating that Treg restrict proliferation of the total activated CD44+ T cell recall response (Fig. 3C; S2C-D). Notably, this increase in Ki67 occurred upon Treg depletion regardless of HSV-2 challenge, in line with prior work with Foxp3^DTR^ mice showing that Treg restrict T cell proliferation at steady state^51^. After HSV-2 challenge, vaginal and dLN CD8 T cells from Treg-depleted and challenged mice also expressed the cytotoxic molecule granzyme B (GrzmB) at a statistically higher frequency and number compared to Treg-sufficient mice, suggesting increased cytotoxic potential (Fig. 3D; S2E-F). Notably, this increase of GrzmB in the vagina and dLN with Treg depletion occurred in the presence or absence of viral challenge, although their frequency was much higher within vaginal tissue (40-60% in VT vs. 5-10% in dLN), and the number of GrzmB+ CD8 T cells in the vagina was statistically higher upon viral challenge in the absence of Treg compared to in the absence of Treg without viral challenge (Fig. S2E-F). We reasoned that the rapid T cell expansion and large increases in GrzmB expression could contribute to the increased tissue damage observed in Treg-depleted and challenged mice (Fig. 2D).

**Figure 3.**
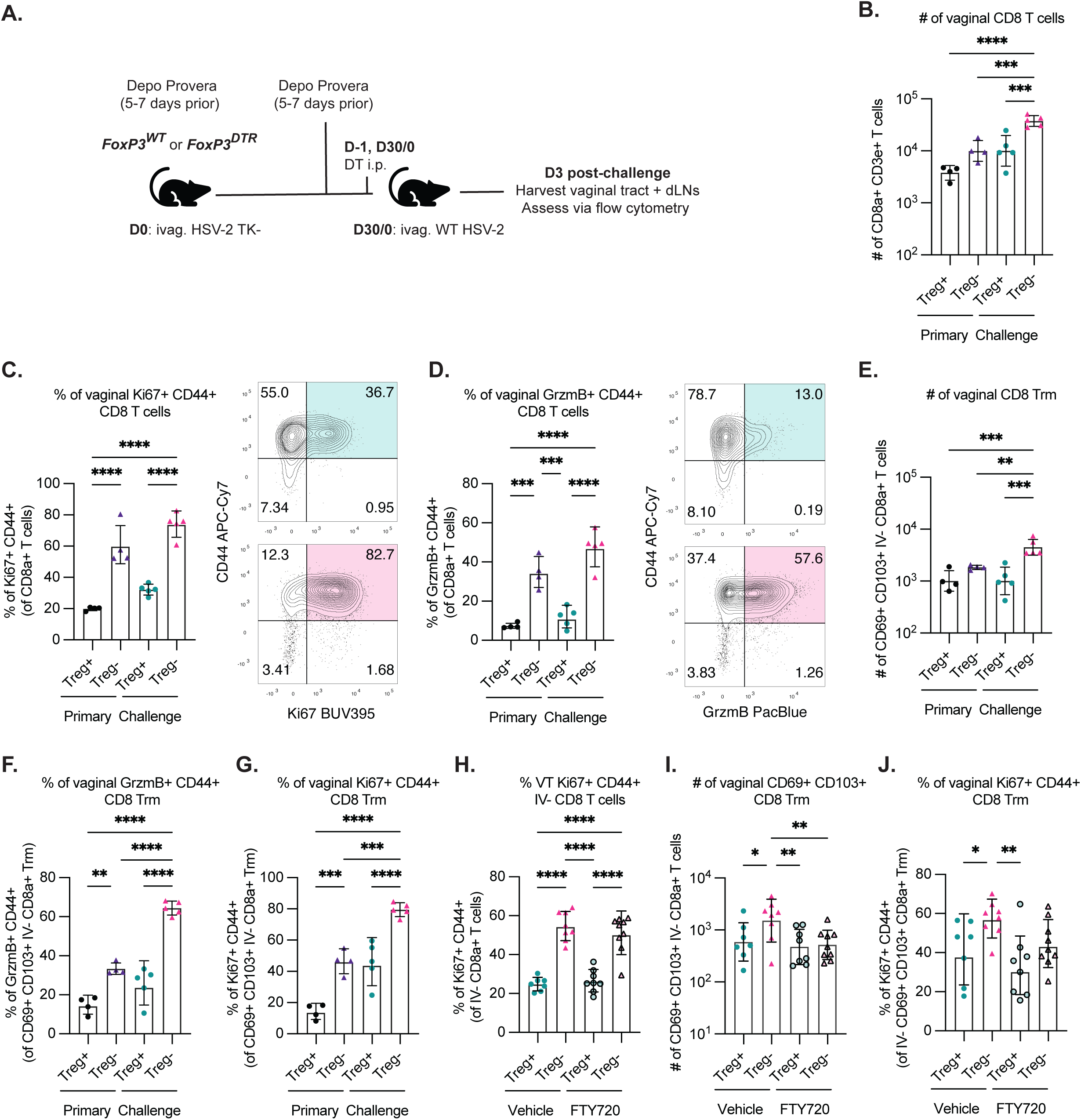
Treg limit the magnitude and cytotoxic potential of the CD8 T cell recall response shortly after WT HSV-2 challenge. (**A**) Schematic shows the experimental design. Female FoxP3^WT^ and FoxP3^DTR^ mice were infected and treated with DT as in Figure 2. VT and dLN were harvested at d3 p.c. and characterized via flow cytometry to assess recall memory CD8 T cell response. (**B**) The absolute number of VT CD8+ T cells in Treg-depleted vs Treg-sufficient unchallenged (“primary”) and challenged (“challenge”) mice. (**C**) Frequency of CD44 and Ki67 expression by VT CD8+ T cells d3 p.c. and representative flow plots (Treg+ challenged group in green, Treg-challenged group in pink). (**D**) Frequency of CD44 and GrzmB expression by VT CD8+ T cells d3 p.c. and representative flow plots (Treg+ challenged group in green, Treg-challenged group in pink). (**E**) Mice were retro-orbitally injected with anti-CD45.2-APC antibody 3 minutes before euthanasia. Trm were defined as CD69+ CD103+ CD45.2 I.V.-label negative. The absolute number of VT Trm. (**F**) Frequency of VT CD44+ GrzmB+ among CD8+ Trm. (**G**) Frequency of VT CD44+ Ki67+ among CD8+ Trm. Data shown are representative of at least two experiments with 4-5 mice per group. Error bars represent the mean and SD. (**H**) Mice were treated with 1mg/kg FTY720 via intraperitoneal injections on days-1, 0, and 2 with respect to WT HSV-2 challenge. Control mice were treated with vehicle (2% β-cyclodextrin in sterile PBS). Frequency of VT Ki67+ CD44+ I.V.-label negative CD8 T cells in FTY720 and vehicle control treated HSV-challenged mice. (**I**) The absolute number of VT CD69+ CD103+ CD8 I.V.-negative Trm in FTY720 and vehicle control treated HSV-challenged mice. (**J**) Frequency of VT Ki67+CD44+ among CD69+ CD103+ CD8 I.V.-negative Trm in FTY720 and vehicle control treated HSV-challenged mice. Data combined from two experiments with 4-5 mice per group. Statistical significance was determined by One-Way ANOVA and Dunn-Šídák multiple comparisons tests in B-J.

Next, we specifically assessed Trm to determine the extent to which Treg may restrict Trm compared to total CD8 T cells found in tissue upon rechallenge. To better identify tissue-resident cells, we injected CD45.2 mice intravenously with anti-CD45.2 APC-conjugated antibody label 3 minutes before euthanasia to label circulating CD45.2+ lymphocytes while sparing those within the tissue^52^ (Fig. S2G). Like the total T cell response, there was a significantly higher number of I.V. label negative (CD45.2-) CD103+ CD69+ CD8 T cells in the VT of Treg-depleted and challenged mice at day 3 p.c. compared to either Treg-sufficient and challenged mice or Treg-depleted and unchallenged mice (Fig. 3E). Additionally, in the absence of Treg, there was a significantly higher frequency and number of vaginal CD44+ Trm expressing Ki67 or GrzmB upon HSV-2 challenge (Fig. 3F-G; S2H-I). Notably, while there was a significant increase in both Ki67 and GrzmB by Trm in the absence of Treg even without HSV-2 challenge, the frequency of Trm expressing each of these markers was further increased upon challenge (Fig. 3F-G), indicating that a combination of Treg removal and infection together boost CD8 Trm proliferation and cytotoxic potential. To assess whether Treg limit proliferation of memory T cells locally during the tissue recall response, we treated mice daily with intraperitoneal (i.p.) injections of FTY720 (an agonist of the S1P receptor that blocks lymphocyte egress from SLOs) starting just before and during challenge. FTY720 administration aimed to prevent T cell egress from the dLN, thus blocking new recruitment to mucosal tissues to determine the effect of Treg on restriction of memory T cell proliferation *in situ*. We found that even upon blockade of T cell recruitment into the vagina during challenge, there was a significant increase in the frequency of I.V.-Ki67+ CD44+ CD8 T cells in vaginal tissue from Treg-depleted compared to Treg-sufficient mice (Fig. 3H). However, their number was not significantly different between FTY720-treated mice irrespective of the Treg presence (Fig. S2J), thus suggesting that though VT Treg limit *in situ* proliferation of CD8 T cells, the increased magnitude of the recall response shortly after challenge in the absence of Treg also relies on the recruitment of effector memory CD8 T cells into the tissue. Similarly, the absolute number of I.V.-CD69+ CD103+ Trm was significantly lower in FTY720-treated Treg-depleted compared to control-treated Treg-depleted mice, as was the number and frequency of Ki67+ CD44+ Trm (Fig. 3I-J; S2K). Altogether, we hereby show that during a tissue recall response, Treg play a critical role in tissue protection by limiting the proliferation and cytotoxic potential of recalled memory T cells within the dLN and vagina, as well as their recruitment and the Trm response at the site of re-infection.

### Treg do not limit the magnitude or cytokine expression of protective HSV-2 gB-speciQc CD8 T cells in the vaginal tissue

Having observed significant increases in the total CD8 T cell and Trm recall response in the absence of Treg, we wanted to determine the impact of Treg depletion on the HSV-specific tissue memory T cell response. The frequency and number of CD44+ CD8 T cells that stained positive with an MHC class I tetramer specific for the HSV glycoprotein B immunodominant epitope (gB) were significantly higher in dLN from Treg-depleted mice regardless of HSV-2 challenge (Fig. S3A-B). In accordance with their numbers, a significantly higher frequency and number of CD44+ gB tetramer+ CD8 T cells expressed Ki67 in dLN from Treg-depleted mice (Fig. S3C). Surprisingly, we did not observe the same significant differences in the frequency or number of CD44+ gB tetramer+ CD8 T cells in the VT following HSV-2 challenge (Fig. 4A-B), and similarly, the frequency and number of HSV-specific CD8 T cells in Treg-depleted and challenged mice that expressed Ki67 compared to Treg-sufficient and challenged mice were comparable (Fig. S3D). Furthermore, there was no difference in the frequency and number of CD69+ CD103+ I.V.-gB tetramer+ CD8 Trm in the VT between Treg-depleted and sufficient groups (Fig. 4C and S3E), nor in the frequency or number of Ki67+ HSV-specific CD8 Trm across the challenged and Treg-depleted groups (Fig. S3F), suggesting that Treg do not limit the expansion of antigen-specific CD8 T cells in the tissue upon re-infection.

**Figure 4.**
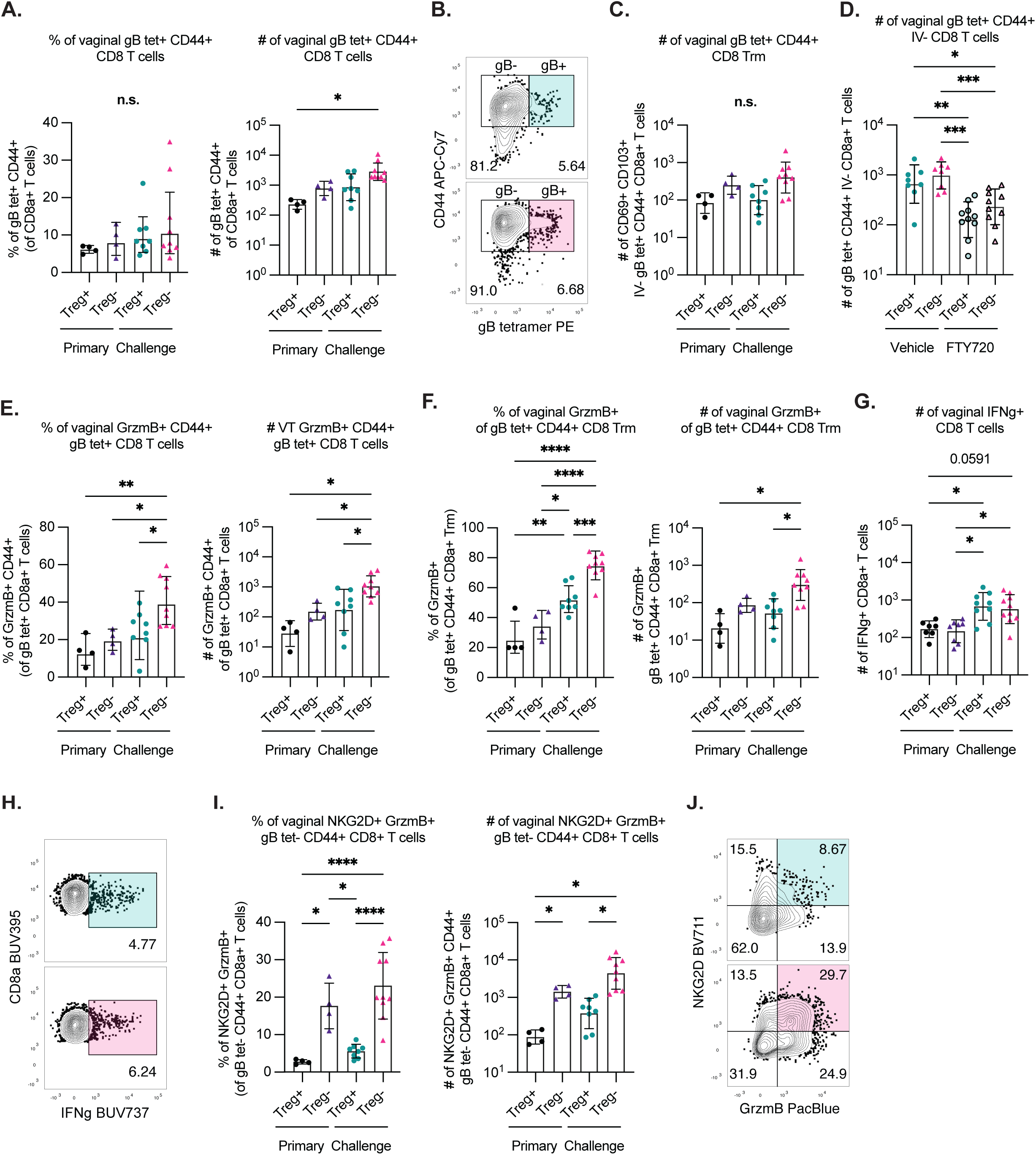
Treg do not limit the magnitude or cytokine expression of protective HSV-2 gB-speciQc CD8 T cells in the vaginal tissue. Female FoxP3^WT^ and FoxP3^DTR^ mice were infected and treated with DT as in Figure 3. (**A**) Frequency (left) and absolute number (right) of gB tetramer+ CD44+ CD8 T cells in VT. (**B**) Representative staining of VT gB tetramer+ CD44+ CD8 T cells and representative flow plots (Treg+ challenged group in green, Treg-challenged group in pink). (**C**) Shown is the absolute number of CD69+ CD103+ gB-tetramer+ I.V.-label negative CD8 T cells in the VT. (**D**) Female FoxP3^WT^ and FoxP3^DTR^ mice were infected and treated with DT as in Figure 3. In addition, mice were treated with 1mg/kg FTY720 via intraperitoneal injections on days-1, 0, and 2 with respect to WT HSV-2 challenge. Control mice were treated with vehicle (2% cyclodextrin in sterile PBS). The absolute number of VT CD69+ CD103+ gB-tetramer+ I.V.-label negative Trm in FTY720 and vehicle control treated HSV-challenged mice. (**E**) Frequency (left) and absolute number (right) of GrzmB+ gB tetramer+ CD44+ T cells. (**F**) Frequency (left) and absolute number (right) of GrzmB+ gB-tetramer+ Trm. (**G**) VT tissue from mice euthanized at 3 days p.c. was processed into a single-cell suspension and stimulated in complete media with gB peptide (immunodominant MHC-I epitope) for 4 hours at 37 °C before conducting intracellular cytokine staining for IFNγ production in CD8 T cells. Shown is the absolute number of VT IFNγ-expressing CD8 T cells.(**H**) Representative intracellular cytokine staining for IFNγ in CD8 T cells (Treg+ challenged group in green, Treg-challenged group in pink). (**I**) Frequency (left) and absolute number (right) of VT NKG2D+ GrzmB+ co-expressing CD44+ gB tetramer negative CD8 T cells (as gated in C). (**J**) Representative staining of NKG2D+ GrzmB+ co-expression in VT CD44+ gB tetramer-negative CD8 T cells (Treg+ challenged group in green, Treg-challenged group in pink). Data shown are combined from two experiments with 4-5 mice per group. Error bars represent the mean and SD. Statistical significance was determined by One-Way ANOVA and Dunn-Šídák multiple comparisons tests in A-G and I.

We next sought to address whether gB-specific CD8 T cells mostly infiltrate from the circulation or expand *in situ* upon challenge. Treating mice with FTY720 just before and during WT challenge revealed that infiltrating gB-specific CD8 T cells comprise a large fraction of the early recall response; forced retention of T cells in the secondary lymphoid organs resulted in a significant decrease of the total absolute number of I.V.-HSV-specific CD8 T cells in the vagina, regardless of the presence or absence of Treg (Fig. 4D). This is consistent with our finding that the frequency of HSV-specific CD8 T cells with a Trm phenotype is lower after challenge compared to at a memory timepoint following primary infection (Fig. S3E), as this likely indicates a large influx of circulating memory CD8 T cells upon challenge that changes the balance of HSV-specific CD8 Trm. The latter could be due at least in part to the known sensing and alarm function of Trm upon cognate antigen recognition in the face of pathogen re-exposure that potentiates the local tissue response with cells from the circulation.

Additionally, we also observed a decrease in the number of Ki67+ I.V.-negative HSV-specific CD8 T cells in FTY720-treated mice, but no significant difference between the challenged groups due to Treg-depletion alone (Fig. S3G). Altogether, these data indicate that while Treg depletion enhances HSV-specific CD8 T cell expansion in the dLN, it does not significantly alter the magnitude or proliferative status of the HSV-specific tissue-resident memory CD8 T cell response in the vagina, which instead seems largely driven by recruitment of circulating effector memory T cells rather than local expansion.

However, just as we observed with the total CD8 T cell and Trm response (Fig. 3), Treg-depleted and HSV-2-challenged mice displayed a significantly higher frequency and number of vaginal HSV-specific CD8 T cells and Trm that were GrzmB+ compared to Treg-sufficient mice or Treg-depleted mice at a memory timepoint following primary infection (Fig. 4E-F; S3H). Notably, the range of frequency for gB tet+ GrzmB+ CD8 T cells in the dLN of challenged mice (∼15-35%) (Fig. S3H) was markedly lower than in the VT (∼40-80%) (Fig. 4E-F). To more deeply characterize CD8 T cell effector function beyond cytotoxic potential, we assessed cytokine expression by T cells following HSV-2 challenge. We stimulated single-cell suspensions of vaginal tissues collected from Treg-depleted and sufficient mice 3 days after HSV challenge with glycoprotein B (SSIEFARL) peptide in complete media for 4 hours at 37°C before intracellular cytokine staining (ICS) for IFNγ. In concordance with our gB tetramer data, there was no difference in the frequency or number of responding IFNγ+ CD8 T cells in the vagina regardless of the presence of Treg, although the number of cytokine-expressing cells was increased following challenge compared to the memory timepoint following primary infection (Fig. 4G-H). Taken together, we demonstrate that Treg allow for the expansion and cytokine effector function of a protective antigen-specific local CD8 T cell response within infected tissues, while limiting the cytotoxic potential of the memory recall response locally, which may selectively minimize cell-mediated tissue damage while still allowing for effective viral control.

Nonetheless, given the comparable magnitude and cytokine expression of antigen-specific responses in vaginal tissue between challenged Treg-depleted and sufficient mice and increased tissue pathology despite clearance of secondary HSV-2 infection, we wondered whether Treg might regulate other memory T cells responding to challenge independent of TCR signaling (i.e., bystander activation). Bystander-activated cytotoxic lymphocytes (BA-CTLs) are transiently activated CD8+ memory T cells that can participate in early anti-pathogen immunity via exposure to cytokines such as type I interferons and pro-inflammatory alarmins, including IL-18, IL-12, and IL-15, rather than via TCR recognition of cognate antigen. BA-CTLs can detect infected cells via NKG2D-dependent recognition of stress ligands and directly kill target cells by delivering cytotoxic granules^13,15^. Our previous work utilizing data from humans with HSV-2 reactivations to inform mathematical modeling^56^ and a murine HSV-2 challenge model after systemic immunization with an irrelevant antigen suggests that BA-CTL recruited to genital tissue contribute significantly to reducing clinical symptoms and early viral burden shortly after viral reactivation or challenge^19^. Thus, we assessed NKG2D and GrzmB co-expression in CD44+ gB tetramer-negative CD8 T cells (Fig. 4I-J; S3I) and found a significantly higher frequency and number within vaginal tissue and dLN from Treg-depleted mice compared with Treg-sufficient mice, although their frequency in Treg-depleted groups was much higher in vaginal tissue (∼20-30%) than dLN (∼5-10%), indicating an overall increase in CD44+ CD8 T cells capable of innate-like recognition via NKG2D and cytotoxic killing in the absence of Treg-mediated restraint.

### Treg restrain the bystander-activated cytotoxic T cell response in mucosal tissue following viral challenge

To formally test whether Treg restrain BA-CTL in the tissue upon challenge, we adoptively transferred 5×10^5^ – 1×10^6^ naïve TCR transgenic Thy1.1+ OT-I CD8 T cells into FoxP3^DTR^ mice one day before primary intravaginal infection with TK-HSV-2-OVA to establish a memory population of OVA-specific CD8 T cells in the vagina. One month later, mice were depleted of Treg and challenged vaginally with WT HSV-2 not expressing OVA (Fig. 5A). As controls, groups of Foxp3^WT^ or Foxp3^DTR^ mice received the adoptive transfer before primary TK-HSV-2-OVA infection, and one month later, mice were treated with DT followed by collection of VT and dLN tissues at a memory timepoint post-primary infection matched to mice that received vaginal HSV-2 challenge (3 days p.c.). As observed in previous experiments (Fig. 4A), we did not and a difference in the number of gB-specific CD8 T cells between Treg-depleted and Treg-sufficient mice (Fig. 5B), suggesting that adoptive transfer of OT-I cells does not alter the magnitude of the recall HSV-specific tissue response. However, following HSV-2 challenge, we observed a significantly higher number of vaginal and dLN Thy1.1+ CD44+ CD8 OT-I cells in Treg-depleted challenged mice compared to all other groups (Fig. 5C; S4A). Additionally, a significantly higher frequency and number of vaginal and dLN Thy1.1+ CD44+ CD8 OT-I cells from Treg-depleted HSV-2-challenged mice were both GrzmB+ and NKG2D+ compared to Treg-sufficient HSV-2-challenged mice or Treg-deficient mice at a memory time point post-primary infection (Fig. 5D; S4B), supporting our hypothesis that Treg restrain the antigen-independent BA-CTL memory response. Vaginal OT-I cells appeared to be more proliferative in HSV-2-challenged than unchallenged mice (Fig. S4C), with a modestly higher number of Ki67+ Thy1.1+ CD44+ OT-I cells in challenged Treg-depleted compared to challenged Treg-sufficient mice (n.s., p = 0.0829). Interestingly, Treg ablation at the time of challenge led to a significantly higher number of bystander-activated NKG2D+ GrzmB+ OT-1 I.V.-Trm (Fig. 5E).

**Figure 5.**
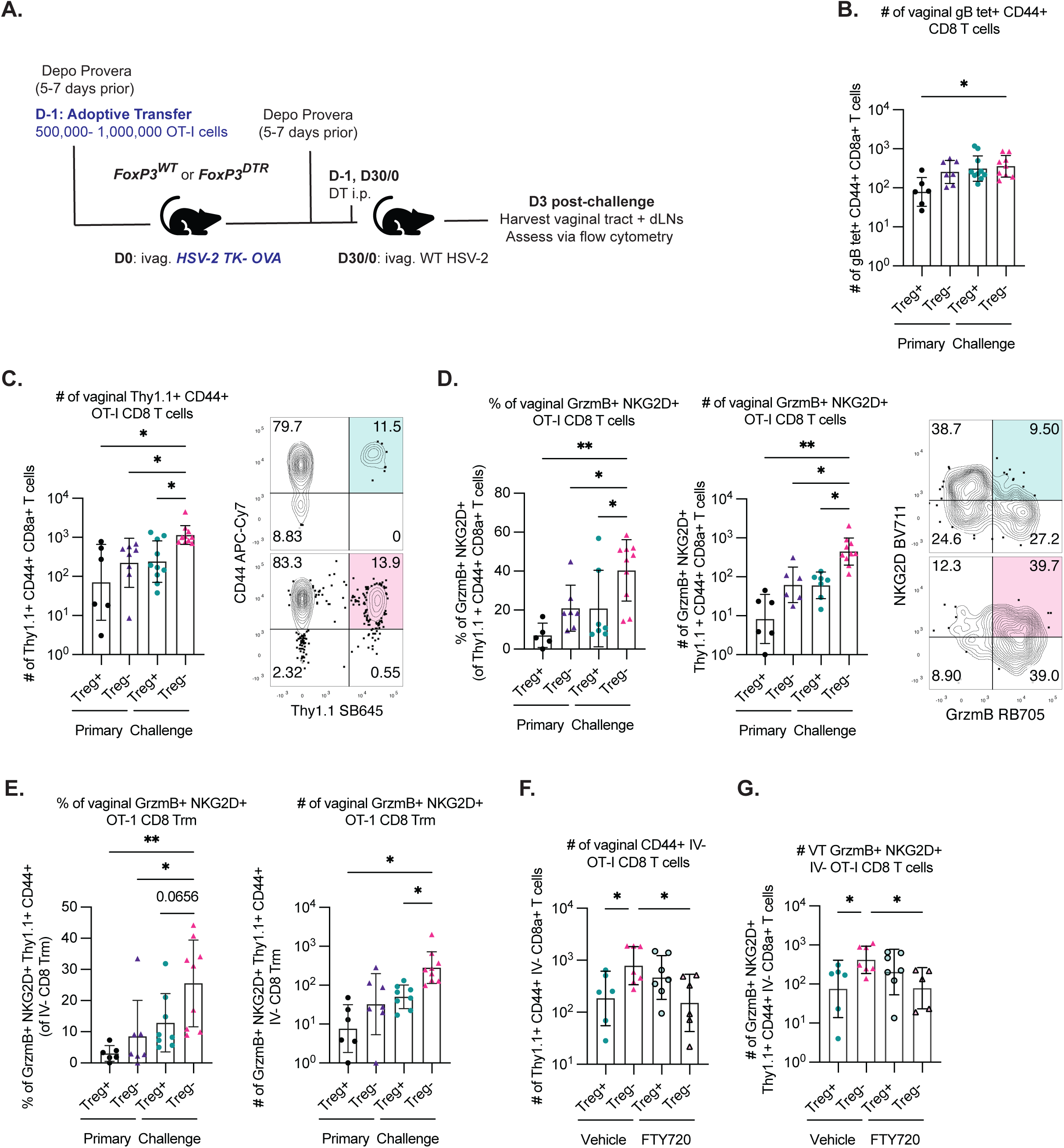
Treg restrain the bystander-activated cytotoxic T cell response in mucosal tissue following viral challenge. (**A**) Schematic shows the experimental design. Thy1.1+ CD45.2+ TCR transgenic OT-I CD8 T cells were first isolated from healthy adult female OT-I mice using an immunomagnetic negative selection kit; 500,000-1 million OT-I cells were then adoptively transferred into female, Depo Provera-treated FoxP3^WT^ and FoxP3^DTR^ mice 24 hours before vaginal infection with 10^5^ PFU of HSV-2 TK-OVA and allowed to recover for 28-30 days. Mice were treated with Depo Provera again 5-7 days before WT HSV-2 (10^4^ PFU) challenge and treated intraperitoneally with diphtheria toxin (DT) to ablate Treg prior to infection (FoxP3^DTR^ mice, Treg-depleted group, Treg-). A dose of 30 μg/kg was administered the day before WT challenge, followed by a dose of 10 μg/kg on d0. VT and dLN were collected d3 p.c. and assessed via flow cytometry. (**B**) Absolute number of VT CD44+ gB tetramer+ CD8 T cells. (**C**) Absolute number of VT CD44+ Thy1.1+ OT-I cells (left) and representative staining (right; Treg+ challenged group in green, Treg-challenged group in pink). (**D**) Frequency and absolute number of VT GrzmB+ NKG2D+ Thy1.1+ CD44+ OT-1 cells (left) and representative staining (right; Treg+ challenged group in green, Treg-challenged group in pink). (**E**) Frequency (left) and absolute number (right) of VT GrzmB+ NKG2D+ Thy1.1+ CD44+ OT-1 Trm (CD69+ CD103+ I.V.-) (**F**) Female FoxP3^WT^ and FoxP3^DTR^ mice were infected and treated with DT as in Figure 3. In addition, mice were treated with 1mg/kg FTY720 via intraperitoneal injections on days-1, 0, and 2 with respect to WT HSV-2 challenge. Control mice were treated with vehicle (2% cyclodextrin in sterile PBS). Shown is the absolute number of I.V.-label-negative VT Thy1.1+ CD44+ OT-1 cells in FTY720 and vehicle-treated control HSV-challenged mice. (**G**) Absolute number of I.V.-label negative VT GrzmB+ NKG2D+ Thy1.1+ CD44+ (bystander-activated) OT-1 cells in FTY720 and vehicle control treated HSV-challenged mice. Data shown are combined from two experiments with 4-5 mice per group. Samples in which the previous gate contained fewer than 25 events were excluded from analysis. Error bars represent the mean and SD. Statistical significance was determined by One-Way ANOVA and Dunn-Šídák multiple comparisons tests in B-G.

As done in previous experiments, we treated mice with FTY720 before and during HSV-2 challenge to determine whether the increase of BA-CTLs in vaginal tissue was primarily due to recruitment or local proliferation. We found that FTY720 treatment significantly decreased the number of total vaginal I.V. label-negative CD44+ OT-I cells and NKG2D+ GrzmB+ CD44+ OT-I BA-CTLs in Treg-depleted mice, suggesting that their increased numbers in untreated Treg-depleted mice are likely due largely to increased recruitment from circulation (Fig. 5F-G).

Thus, our findings are consistent with a role for Treg in limiting the magnitude of the BA-CTL tissue response upon secondary intravaginal infection, thereby coordinating an appropriate, pathogen-specific memory T cell response that is finely regulated to clear the pathogen while limiting cytotoxic activity that could compromise host tissue integrity.

### Treg limit the cytotoxicity of antigen-speciQc and bystander CD8 T cells via an IL-2-dependent mechanism

Next, we wanted to further investigate mechanisms by which Treg can suppress the expansion and cytotoxic potential of memory T cells, and particularly the BA-CTL. A canonical mechanism of Treg immunosuppressive function is their ability to sink IL-2 availability in the local immune environment through expression of the high-affinity IL-2 receptor subunit, CD25^23^. Their efficient binding and obligatory consumption of IL-2 sequesters it from other T cells, depriving them of the potent activating cytokine. Furthermore, others have shown that strong IL-2 signaling is necessary to induce cytolytic function in effector CD8 T cells^57^, and more recently, it was shown that IL-2 was able to modestly induce expression of Granzyme B and NKG2D by human memory CD8 T cells, though to a lesser extent compared to IL-15^58^.

Thus, we hypothesized that in the absence of Treg, there is more IL-2 available in the local immune milieu, resulting in the activation and expansion of CD8 T cells leveraged in the memory tissue recall response.

To address our hypothesis that unrestrained availability of IL-2 in the absence of Treg was contributing to the expansion and activation of BA-CTL and other effector T cells in our HSV-2 challenge model, we blocked IL-2 signaling *in vivo* prior to WT challenge using a monoclonal anti-IL-2 antibody (clone JES6-1A12)^59^ (Fig. 6A). IL-2 blocking just before challenge was associated with a significantly lower frequency and number of total GrzmB+ CD44+ CD8 T cells within the vagina and dLN of Treg-depleted mice compared to isotype-treated Treg-depleted mice also challenged with HSV-2 (Fig. 6B; S4D). IL-2 blockade did not impact the magnitude of the HSV gB-specific CD8 T cell response within the vagina and dLN (Fig. S4E). Additionally, there was no significant difference in the frequency or number of vaginal HSV-specific CD8 T cells expressing GrzmB from Treg-depleted mice upon neutralization of IL-2 (Fig. 6C), though in the dLN, IL-2 was required for the increase observed in the frequency of HSV-specific CD8 T cells expressing GrzmB (Fig. S4F). Further, just as in our initial experiments (Fig. S3), there was no significant difference in the frequency of Ki67 expression by vaginal HSV-specific CD8 T cells between Treg-sufficient and depleted challenged mice treated with anti-IL-2 or isotype control monoclonal antibodies (Fig. 6D; S4G). Notably, IL-2 blocking significantly inhibited the frequency and number of total CD44+ gB tetramer-negative NKG2D+ GrzmB+ vaginal CD8 T cells (Fig. 6E), and specifically bystander activation of vaginal and dLN Thy1.1+ cells (as identified by co-expression of NKG2D and GrzmB) and their expression of Ki67 in Treg-depleted mice (Fig. 6F-G; S4H). Altogether, our findings suggest that Treg limit available IL-2 in the immune environment to restrain bystander activation, cytotoxic potential, recruitment, and proliferation of memory CD8 T cells during a tissue recall response.

**Figure 6.**
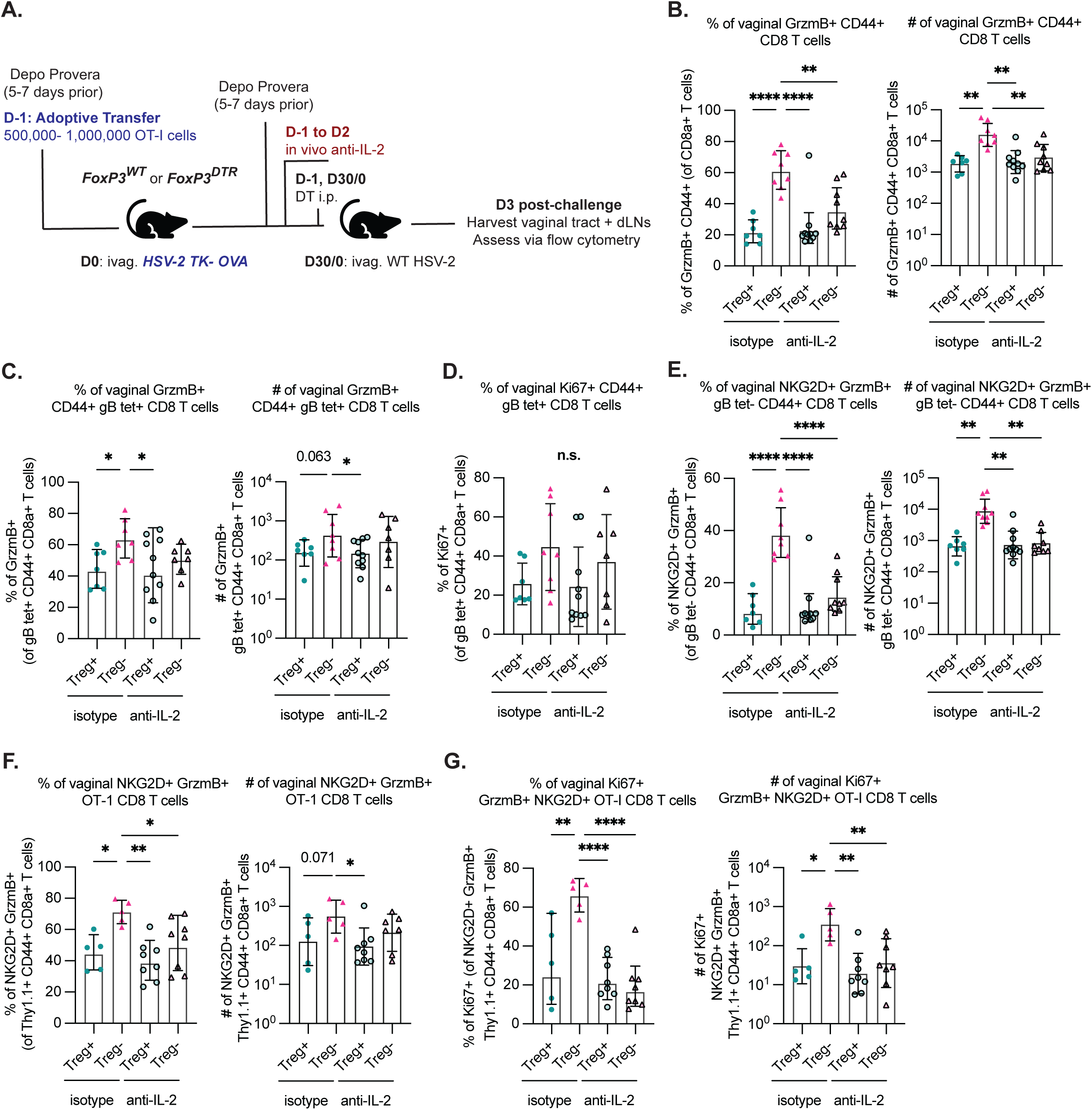
Treg limit the cytotoxicity of antigen-speciQc and bystander CD8 T cells via an IL-2-dependent mechanism. (**A**) Schematic shows the experimental design. 500,000-1 million OT-I cells were adoptively transferred into female Depo-Provera-treated FoxP3^WT^ and FoxP3^DTR^ mice 24 hours before vaginal infection with 10^5^ PFU of HSV-2 TK-OVA and allowed to recover for 28-30 days. Mice were again treated with Depo Provera 5-7 days before WT HSV-2 (10^4^ PFU) challenge and treated intraperitoneally with diphtheria toxin (DT) to ablate Treg prior to infection (FoxP3^DTR^ mice, Treg-depleted group, Treg-). Mice were also administered both intravenous (150µg) and intravaginal (80µg) anti-IL-2 monoclonal antibody (clone JES6-1A12) or rat IgG2a isotype control on day-1 with respect to WT challenge. 40µg of anti-IL-2 or isotype control was administered intravaginally d1 and 2. VT and dLN were collected 3 days p.c. and assessed via flow cytometry. (**B**) Frequency (left) and absolute number (right) of total VT CD44+ GrzmB+ CD8 T cell response in WT HSV-2 challenged Treg-depleted or sufficient mice after treatment with anti-IL-2 or isotype control. (**C**) Frequency (left) and absolute number (right) of GrzmB expression in VT gB tetramer+ CD44+ CD8 T cells from WT HSV-2 challenged Treg-depleted or sufficient mice after treatment with anti-IL-2 or isotype control. (**D**) Frequency of Ki67+ in VT gB tetramer+ CD44+ CD8 T cells from WT HSV-2 challenged Treg-depleted or sufficient mice after treatment with anti-IL-2 or isotype control. (**E**) Frequency (left) and number (right) of VT NKG2D+ GrzmB+ CD44+ gB-tetramer negative CD8 T cells. (**F**) Frequency (left) and absolute number (right) of NKG2D+ GrzmB+ OT-I BA-CTL in VT from WT HSV-2 challenged Treg-depleted or sufficient mice after treatment with anti-IL-2 or isotype control. (**G**) Frequency of Ki67 expression in VT NKG2D+ GrzmB+ OT-I BA-CTL from WT HSV-2 challenged Treg-depleted or sufficient mice after treatment with anti-IL-2 or isotype control. Data shown are combined from two experiments with 3-5 mice per group. Error bars represent the mean and SD. Statistical significance was determined by One-Way ANOVA and Dunn-Šídák multiple comparisons tests in B-G.

### Antigen-presenting cells increase IL-15 trans-presentation in response to available IL-2 in the absence of Treg

In addition to depriving memory CD8 T cells of IL-2, we hypothesized that Treg might additionally restrain the recall tissue response via regulation of IL-15 trans-presentation by antigen-presenting cells (APC) to responding T cells. It is well known that IL-15 is an essential cytokine that can induce cytotoxicity in memory CD8 T cells, even in the absence of TCR specificity and antigen recognition^12,16–20,58,60–62^. IL-15 must be trans-presented by IL-15Ra to be biologically active and is constitutively expressed by many different cell types (including monocytes, macrophages, and dendritic cells^63^), and can be further elevated by type I IFN or pathogen recognition receptors in the case of viral infection^64–66^. We previously demonstrated that Treg limit IL-15 trans-presentation in the context of primary flavivirus infections^50,67^. Thus, we assessed IL-15 trans-presentation by APC upon Treg depletion and HSV-2 challenge via flow cytometry and found a significantly higher number of IL-15+ myeloid APC (Ly6G-CD19-CD3e-CD45+) in vaginal tissue from Treg-depleted mice HSV-2-challenged compared to Treg-sufficient HSV-2-challenged mice, as well as compared to Treg-depleted mice at a memory timepoint following primary infection (Fig. 7A; S5A-C). This indicates that both HSV-2 challenge and removal of Treg-mediated restraint together contribute to an increase in IL-15 trans-presentation by myeloid APC. Although Treg-depleted and challenged mice had a significantly higher number of IL-15+ APC in dLN compared to Treg-sufficient mice, Treg-depleted unchallenged mice also had elevated numbers (Fig. 7A; S5C). Notably, the majority of IL-15+ APC were composed of monocytes, followed by macrophages (Fig. 7A). We wondered if an increase in available IL-2 in the absence of Treg might impact APC activation and IL-15 trans-presentation, either indirectly through CD4 Tconv interactions or directly via APC IL-2-receptor (IL-2R) signaling. Activated monocytes and macrophages have been shown to express a functional IL-2R that, upon IL-2 binding, promotes anti-microbial functions such as intracellular killing of bacteria, tumoricidal activity, and IFNγ production^68–72^. To determine whether there was a relationship between IL-2 levels and IL-15 trans-presentation, we evaluated IL-15+ APC in challenged Treg-sufficient and depleted mice after in vivo IL-2 blocking. Indeed, we found that IL-2 blockade results in decreased frequencies of IL-15 trans-presentation by macrophages, monocytes, and dendritic cells, and specifically a statistically significant decrease in the number of IL-15+ macrophages within vaginal tissue from HSV-2-challenged Treg-depleted mice (Fig. 7B-E; S5D-F). Conversely, IL-2 blockade had less of an effect on the frequency and number of IL-15 trans-presentation by myeloid cells in dLN (Fig. S5G-L).

**Figure 7.**
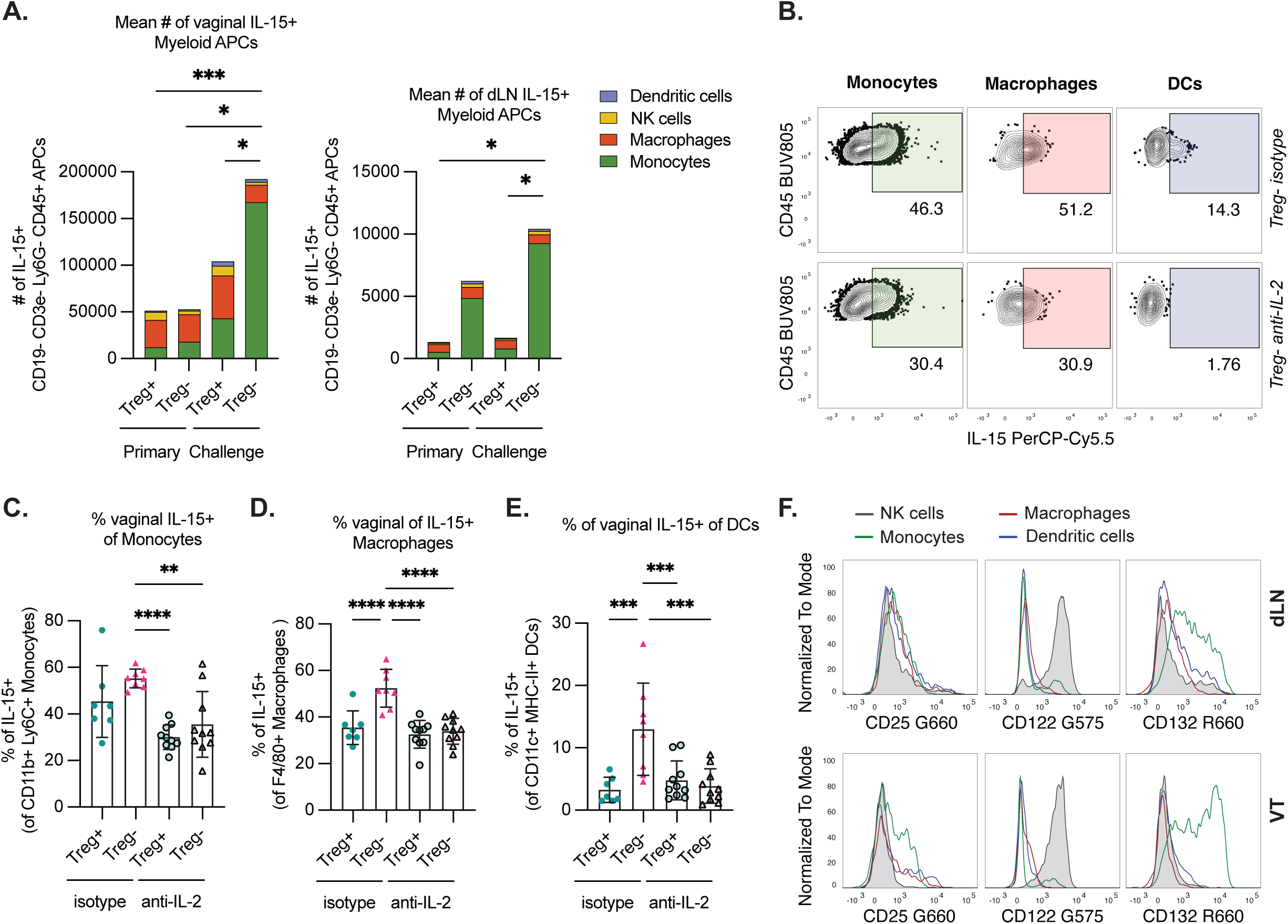
Antigen-presenting cells increase IL-15 trans-presentation in response to available IL-2 in the absence of Treg. (**A**) The absolute number of IL-15+ CD19-CD3e-CD45+ APC is shown. Stacked bar plot displays cell type subset as a fraction of the total mean number of IL-15+ myeloid APC in the VT (left) and dLN (right) at day 3 p.c. Data shown are representative of two experiments. (**B**) Representative flow staining of total IL-15 expression in monocytes, macrophages, and DCs from Treg-depleted and challenged mice treated with isotype control (top) or anti-IL-2 (bottom) Mab. Female FoxP3^WT^ and FoxP3^DTR^ mice were infected and treated with DT as in Figure 1, and with anti-IL-2 or isotype control Mab as in Figure 5. (**C**) VT and dLN were collected 3 days p.c. and assessed via flow cytometry. Shown is the frequency of IL-15 trans-presentation by monocytes (CD3e-CD19-Ly6G-), (**D**) macrophages (CD3e-CD19-Ly6G-NK1.1-F4/80+), and (**E**) DCs (CD3e-CD19-Ly6G-NK1.1-F4/80-CD11c+ MHC-II+) in VT from WT HSV-2 challenged (day 3 p.c.) Treg-depleted or sufficient mice after treatment with anti-IL-2 or isotype control Mab. Data shown are combined from two experiments with 4-5 mice per group. Error bars represent the mean and SD. Statistical significance was determined by One-Way ANOVA and Dunn-Šídák multiple comparisons tests. (**F**) Representative histograms showing expression of IL-2R components (CD25, CD122, CD132) by NK cells (CD3e-CD19-Ly6G-NK1.1+) used as an IL-2R+ control, monocytes (CD3e-CD19-Ly6G-CD11b+Ly6C+), macrophages, and dendritic cells from VT tissues of unchallenged Treg-sufficient mice about a month after primary TK-infection (normalized to mode).

To ascertain whether increased IL-15 trans-presentation in the absence of Treg could be in part a direct effect of sensing of IL-2 by APC, we first evaluated expression of the IL-2R components *in vivo* by APC in vaginal tissue and dLN of Treg-sufficient unchallenged mice one month after primary intravaginal HSV-2 infection and found that monocytes, macrophages, and dendritic cells all express CD25 and CD132 to equal or higher levels than observed in NK cells (Fig. 7F; S5M). While CD122 was expressed at lower levels in comparison to NK cells, a fraction of monocytes in particular stained positive for CD122 (Fig. 7F). To further assess the relationship between available IL-2 levels and IL-15 trans-presentation *in vitro*, splenocytes were isolated from healthy naïve FoxP3^WT^ mice; half of the single cell suspension was directly plated at 1 million cells/well, while the remaining half was put through a CD3e T cell magnetic microbead depletion kit before plating at 1 million cells/well. The total vs. CD3e-depleted splenocytes were then cultured with increasing concentrations of recombinant murine IL-2 *in vitro* overnight at 37C°. We found that only macrophages directly responded to IL-2 stimulation with significant increases in the frequency of IL-15 trans-presentation, in a dose-dependent manner, in CD3e-depleted cultures (Fig. 8A-B). Additionally, in IL-15+ macrophages, we observed a similar dose-dependent increase in the frequency of CD25 expression (Fig. 8C). We did not observe the same response to increasing IL-2 in monocytes and dendritic cells *in vitro* (Fig. 8D-E). To our knowledge, this is the first report of high levels of IL-2 directly increasing IL-15 trans-presentation in macrophages and adds partial mechanistic insight into the pathways by which Treg coordinate the recalled cytotoxic response upon secondary infection.

**Figure 8.**
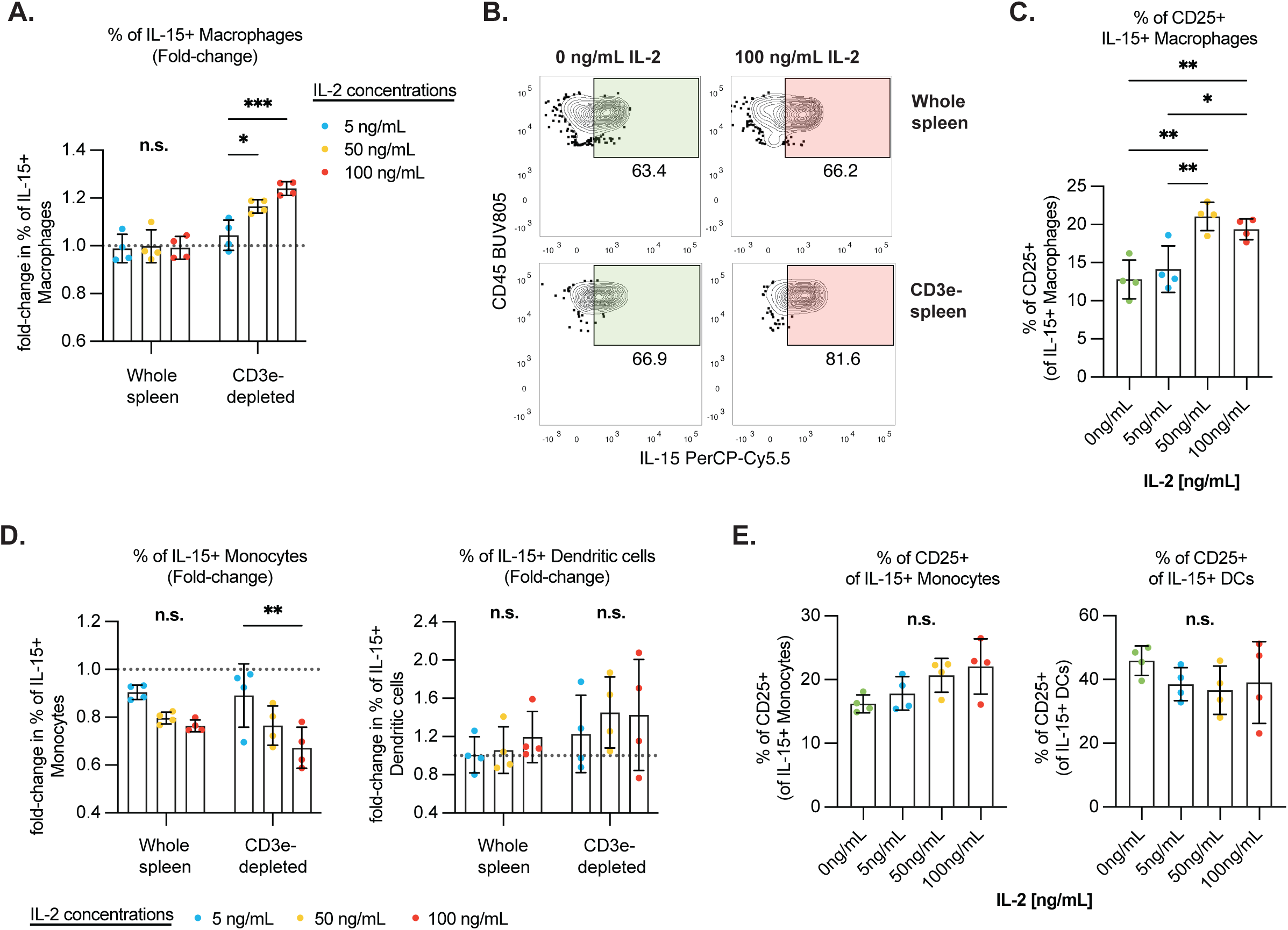
Macrophages upregulate IL-15 trans-presentation in response to increasing IL-2 concentrations *in vitro*. Single-cell suspensions of total spleen and CD3e-depleted spleen (isolated via CD3e T cell magnetic microbead depletion kit) from healthy FoxP3^WT^ mice were plated at 1 million cells/well and stimulated with increasing concentrations of recombinant murine IL-2 (rmIL-2) in complete media overnight at 37 °C. (**A**) Frequency of IL-15 trans-presentation by macrophages after overnight culture with increasing concentrations of rmIL-2. Data presented as a fold-change in frequency in comparison to the media-only control (0ng/mL of rmIL-2). (**B**) Representative flow plot of IL-15+ gating of macrophages (from total vs. CD3e-depleted spleen). (**C**) Frequency of extracellular CD25 expression by IL-15+ macrophages within CD3e-depleted splenocyte samples after overnight rmIL-2 culture. (**D**) Frequency of IL-15 trans-presentation by monocytes (left) and DCs (right) after overnight culture with increasing concentrations of rmIL-2. Data presented as a fold-change in frequency in comparison to the media-only control (0ng/mL of rmIL-2). (**E**) Frequency of extracellular CD25 expression by IL-15+ monocytes (left) and DCs (right) within CD3e-depleted splenocyte samples after overnight rmIL-2 culture. Data shown are representative of two experiments using splenocytes from 4 individual FoxP3^WT^ mice. Error bars represent the mean and SD. Statistical significance was determined by Two-Way ANOVA and Tukey’s multiple comparisons tests for A and D, or One-Way ANOVA and Dunn-Šídák multiple comparisons tests for C and E.

## DISCUSSION

Mucosal barrier tissue sites are continuously exposed to a mix of pathogens and environmental antigens, including commensals. As the most abundant memory T cell subset, Trm are critically important to coordinating rapid and long-term immunity against infections at mucosal surfaces^7,8,73^. Despite the sustained interest and research efforts in understanding their formation, activation, and localization since their discovery over 2 decades ago^3,4,7,74,75^, we still lack a clear understanding of Trm regulation, particularly during a tissue recall response, which motivated our study of Treg. We reasoned that regulation of memory recall responses would be critical upon re-infection, given that an over-exuberant tissue recall response could lead to significant tissue injury, thereby contributing to disease, particularly in chronic or episodic, frequently recurrent infections such as HSV-2^76^. Treg have been shown to infiltrate and persist alongside HSV-2-specific Trm throughout lesion healing^46,77^, though their role in shaping Trm recall responses upon reactivation remains unclear. Others have reported the localization and specialization of non-lymphoid tissue Treg in the context of inflammatory settings, such as viral respiratory infection, sterile muscle damage, and adipose-tissue inflammation^78–84^. Furthermore, though it is well known that Treg limit excessive inflammatory T cell responses in a myriad of direct and indirect mechanisms, how and whether Treg modulate recalled memory T cells and non-specific BA-CTL responses to infection is not well understood.

We and others have shown that human and murine Treg in non-lymphoid tissues have a more activated and immunosuppressive transcriptional and phenotypic signature compared to Treg in circulation and lymphoid tissue^21,44,45,80,81,84–86^. Given this heightened activation, in addition to their presence in barrier tissues, we hypothesized that they participate in anti-pathogen immunity upon mucosal infection and re-infection. It is plausible that Treg also possess the ability to form immunological memory just as conventional T cells do. In support of this idea, previous reports have documented the formation of memory Treg^87–91^. A study from the Rudensky group revealed that prior exposure to an autoinflammatory environment or systemic LCMV infection imprints a stable non-lymphoid tissue localization preference for Treg, as well as a limited set of transcriptional changes that are consistent with a memory of prior inflammation^78^. However, they also observed that Treg reversed many, though not all, activation-induced changes. Many studies assessing tissue Treg or memory Treg are carried out in settings of autoimmunity, steady-state, systemic infection, or sterile tissue inflammation or injury^78,79,82,90^. Yet, such models do not precisely mimic natural pathogen encounter or chronic exposure, in which microbial antigens and pathogen-associated molecular patterns drive distinct inflammation types and responses in addition to tissue-specific changes in the microenvironment due to localized pathogen replication. Therefore, we hypothesize that Treg form memory responses to mucosal virus infection, allowing them to be poised to respond to re-infection within affected tissues.

Here, using a murine model of localized mucosal viral infection, we demonstrate that Treg limit the expansion and cytotoxic potential of the recalled memory CD8+ T cell and non-specific BA-CTL response to mucosal viral challenge by sinking local IL-2 and sequestering it from conventional T cells and APC. We have previously demonstrated that at steady state and in the context of primary infection, Treg limit IL-15 trans-presentation by antigen-presenting cells, including DCs and monocytes^50,92^. Here, we build on that work to now focus on the tissue during a recalled memory response. Further, we propose that restricted availability of IL-2 in the local immune environment in the presence of Treg restrains T cell activation, proliferation, and survival, plus limits trans-presentation of IL-15 by APC to T cells. Previous work identified IL-15 as a major player in the induction of the cytotoxic program in CD8 T cells^20,58,93^, yet regulatory mechanisms of local IL-15 within tissues remained unclear. Mucosal tissues, including the genital tract, are frequently exposed to inflammatory threats and commensals that could induce IL-15, suggesting that this response must be subject to immunoregulation to restrict the recall memory CD8 T cell response within tissues and avoid immunopathology.

Notably, while Lee *et al.* investigated cell-intrinsic mechanisms of restraint of the IL-15-induced bystander activation of memory CD8 T cells and found that TCR signaling could quell the cytokine-induced cytotoxic program^58^, this does not account for regulation of cytokine-induced cytotoxicity in the absence of cognate antigen. The latter is predicted to occur frequently given the diverse range of microbial exposures encountered at mucosal barrier tissue sites. Thus, our work for the first time focuses on extrinsic environmental and cellular cues in mucosal tissues that restrain recall memory CD8 T cell cytotoxic activity, and notably BA-CTL. Though IL-2 and IL-15 signaling share the common gamma chain (CD132) and IL-2/IL-15 receptor beta chain (CD122), they seem to play different roles in immunity. Notably, IL-15 is produced by many cell subsets, including non-myeloid epithelial and stromal cells. IL-15 expression can be induced both in response to microbial pathogen-associated triggers^94^ and in instances of sterile inflammation like autoimmunity^95^. IL-15 overexpression is commonly linked to various tissue-specific autoimmune disorders such as rheumatoid arthritis, multiple sclerosis, systemic lupus erythematosus, celiac disease and insulin-dependent diabetes^96–99^. Interestingly, in the context of cancer, metastatic patients with low expression of IL-15 due to a deletion at the IL-15 genetic locus have an increased risk of relapse and lower density of proliferating intratumoral immune cells, hinting at a role for IL-15 in tissue immunity and pathology^100,101^. In contrast, IL-2 deficiency as opposed to overexpression typically contributes to autoimmunity that can be ameliorated by IL-2-based therapies that target *in vivo* expansion of Treg to restore immune homeostasis^102,103^. Our findings here support a model wherein the tissue microenvironment keeps memory recall responses within tissues in check, at least in part through Treg control of available IL-2 and IL-15.

Finally, we propose that Treg, in addition to their direct modulation of both innate and adaptive responses, are also critical in preventing sustained excessive local inflammation that contributes to pathologic BA-CTL responses. Importantly, Treg restrain the cytotoxic potential of the total memory recall T cell responses and immune-mediated tissue damage upon secondary mucosal viral infection, while still allowing for an optimal pathogen-specific protective tissue T cell response. The fact that the presence or absence of Treg did not significantly impact the immunodominant HSV-2-specific CD8 T cell tissue memory recall response in terms of size and cytokine expression is particularly interesting. A growing body of research on the role of Treg during infection supports the idea that they are essential in limiting excessive inflammation that can distract the immune response from focusing on the formation of protective pathogen-specific T cell responses^43,44,87,92,104^. Our research group has previously shown that Treg are a critical source of TGF-β within infected tissues, shaping the generation of CD103+ CD8+ Trm and supporting their maintenance in brain tissue after primary WNV infection in mice^92^. Furthermore, recent work by Jarjour et al. shows that, once formed, CD8+ memory T cells can dynamically adapt to changes in the cytokine milieu during viral infection, making them resilient to potential loss of homeostatic signals^105^. Taken together, by modulating the cytokine environment, Treg are critically important in coordinating priming of classical, Ag-specific T cells and Trm formation within tissues; yet once differentiation establishes a Trm program in T cells, their survival may be less impacted by cytokine alterations upon Treg-depletion during infection and even supported by compensatory inflammatory cytokines.

Here, we have demonstrated a role for Treg in directing the recalled memory CD8 T cell response to viral re-challenge (Fig. 9). Importantly, Treg soak up excess IL-2 in the dLN and in the local tissue microenvironment, thereby depriving memory CD8 T cells of this cytokine but also restricting local IL-15 trans-presentation by APC. Treg-mediated restriction of IL-2 in both tissue sites limits overall CD8 T cell proliferation, activation, and induction of the cytotoxic program, although the range of frequencies of GrzmB expression by CD8 T cells in Treg-depleted mice from the dLN was much lower than those observed in the vagina and likely due to less innate signaling in the dLN compared to the infection site that could contribute to induction of IL-15 transpresentation (i.e., Type I IFN and pathogen-associated molecules).

**Figure 9.**
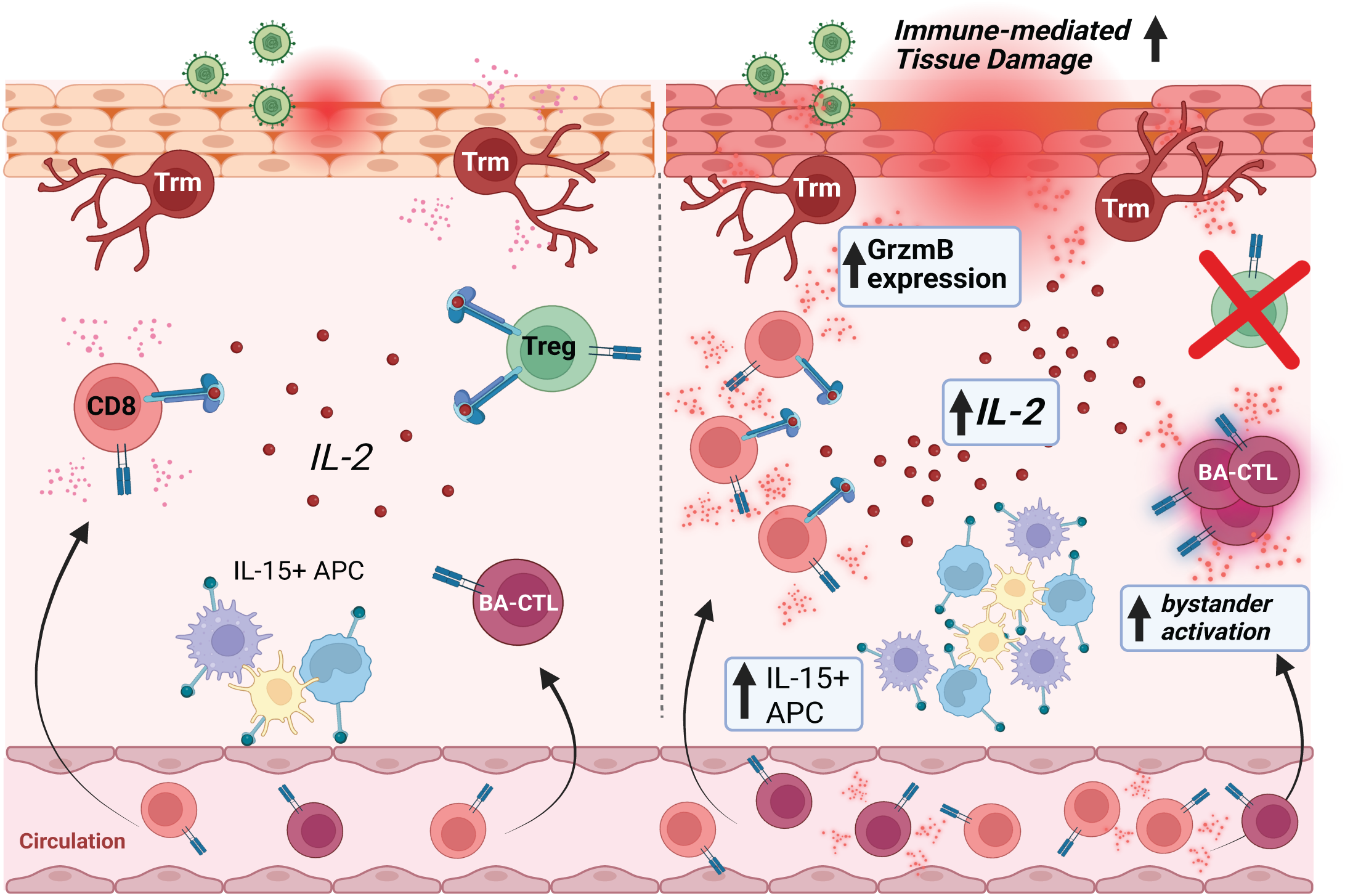
**Hypothesized framework for how regulatory T cells restrain cytotoxic and bystander CD8 T cells at least in part through IL-2/IL-15 mediated mechanisms. LEFT**: Viral challenge in the presence of intact and functional Treg allows for the expansion and recruitment of memory CD8 T cells (both viral antigen-specific and non-specific) to the site of exposure, the mucosal barrier tissue. Upon challenge, antigen-presenting cells upregulate IL-15 trans-presentation in response to increased IL-2, type I IFN, and sensing of pathogen-associated molecules. Bystander-activated memory CD8 T cells present in the tissue are transiently activated by increased IL-15 and type I interferons and contribute to controlling the spread of the virus shortly after infection^19^, bridging the innate and adaptive immune response. **RIGHT**: Viral challenge in the absence of Treg results in overall increased activation, expansion, and recruitment of memory CD8 T cells into the tissue. With increased numbers of recalled memory T cells, available IL-2 levels increase locally without Treg present to sequester and consume excess. Increased activation of memory CD8 T cells within the mucosal tissue leads to increased frequencies and numbers of IL-15-transpresenting antigen-presenting cells locally (macrophages directly upregulate IL-15 in response to IL-2, whereas monocytes and dendritic cells appear to do so indirectly via other signals/mechanisms). Though Treg depletion does not impact the magnitude of the pathogen-specific CD8 T cell response, the large increase in available IL-15 and IL-2 contributes to increased expansion and recruitment of bystander-activated cytotoxic T cells and overall granzyme B expression in all CD8 T cells, resulting in increased immune-mediated tissue damage locally and worsened disease outcomes despite proper clearance of infection.

Thus, we demonstrate for the first time that Treg govern environmental cues that induce the cytotoxic program of memory CD8 T cells, including bystander activation. Importantly, Tregs do not restrict the protective, antigen-driven and antigen-specific tissue memory CD8 T cell response, as demonstrated by the lack of difference in tissue pathogen control, though they are critical to reducing tissue injury in the context of a tissue recall response. Altogether, our findings reveal the importance of Treg as critical mediators of innate and adaptive recall responses within mucosal tissues, with implications for the design of successful vaccines that elicit protective, but not pathogenic, Trm populations poised to respond at mucosal sites of pathogen exposure and entry.

### Limitations of the study

We assessed bystander activation using an adoptive transfer model that leveraged OT-I cells to ensure that cells were not activated via TCR signals (Fig 5). We recognize that assessment of endogenous rather than TCR transgenic T cells would be preferable. We did attempt to establish an endogenous OVA-specific memory CD8 T cell response in the mucosa by priming mice intravaginally with TK-HSV-2-OVA and then assessing the frequency of endogenous GrzmB+ NKG2D+ OVA-specific T cells after depletion of Treg and WT HSV-2 challenge. Unfortunately, this strategy did not establish sufficient numbers of OVA-specific memory CD8 T cells in the VT to properly enumerate and assess their bystander activation phenotype after challenge (generally, less than 25 endogenous OVA-specific CD8 T cells could be detected in the vagina). Thus, we instead relied on adoptive transfer to increase the number of T cells detectable in the vagina.

Additionally, although we report that increased IL-2 availability results in a direct dose-dependent increase in IL-15 trans-presentation by macrophages, we have not addressed the exact mechanism by which IL-2R signaling results in this response from APC. Future studies aimed at understanding the distinct effects of IL-2 and IL-15 on immune cells present in non-lymphoid tissue would aid in the improved design of vaccines and therapeutics to achieve improved immunity and minimize collateral tissue damage.

## METHODS

### MICE

6–10-week-old female wildtype C57BL/6J (Jackson Laboratories, Bar Harbor, ME, strain #000664), FoxP3 ^eGFP^ (C.Cg-Foxp3tm2Tch/J), or B6.129(Cg)-Foxp3tm3(HBEGF/GFP) Ayr “FoxP3 ^DTR^” knock-in mice (a gift from Dr. Alexander Rudensky, bred at Fred Hutch) were used for in vivo experiments and in vitro stimulations. C57BL/6-Tg (Tcra/Tcrb)1100Mjb/J OT-I Thy1.1 were used for isolation of CD8 OT-I from splenocytes for adoptive transfers. All animal experiments were approved by Fred Hutch IACUC, and the study was conducted in strict compliance with the PHS Policy on Humane Care and Use of Laboratory Animals. Due to the HSV-2 infection route used in this model (vaginal), only female mice were used.

### INFECTIONS

Mice were injected subcutaneously in the neck ruff with 2mg of medroxyprogesterone acetate (Depo-Provera) dissolved in sterile PBS, 5–7 days prior to intravaginal (ivag.) infection. Mice were infected with either 10^5^ PFU TK-HSV-2 (HSV-2 TK-/kpn) or 10^5^ PFU of TK-OVA HSV-2 (HSV-2 OVA TK-/kpn) for primary infection. Mice were challenged 30 days post-primary HSV-2 TK-infection with 10^4^ PFU of wild-type HSV-2 (186syn+) derived from a human clinical isolate. Unchallenged/uninfected mice were also administered Depo-Provera to control for any estrus cycle effects on immunity.

### INTRAVASCULAR LABELING

Mice were administered an intravenous (i.v) retro-orbital injection (ROI) of APC-CD45.2 (3µg in 200µL per mouse) three minutes prior to CO^2^ euthanasia to label and differentiate circulating lymphocytes from tissue-resident cells.

### MOUSE TISSUE PROCESSING

Murine vaginal tracts (VT), including the cervix, were harvested into RP10 media and minced with scissors prior to a 30 min incubation in DM10 (10% heat-inactivated FBS, 2 mM L-glutamine, 100 U/mL penicillin-streptomycin, 5 mM sodium pyruvate) containing collagenase D (2mg/mL; Sigma Cat.11088858001) and DNAse I (15µg/mL; Sigma Cat.10104159001) at 37°C with gentle shaking. Tissue samples were then mashed through a 100um strainer and washed with 20mL of RP10 media. VT samples were spun down, and the entire cell pellet was resuspended with fluorescence-activated cell sorting (FACS) buffer into a single-cell suspension stained and used for flow cytometric analysis. Vaginal-draining lymph nodes (dLN), iliac and inguinal, were harvested into RP10, mashed through a 100um strainer and resuspended in FACS buffer to prepare a single-cell suspension. Spleens were harvested into RP10, mashed through a 100um strainer, and incubated in 5 mL ACK lysis buffer for 5min. The ACK-treated cells were suspended in RP10 medium before proceeding with flow staining. All samples were spun down at 300g for 5 min to pellet. All samples were put through 35µm nylon strainer shortly before flow cytometric analysis.

### FLOW CYTOMETRY

Cells were incubated in fixable Live/Dead amine-reactive blue viability dye (Thermo Fisher Cat. L23105) for 25 min (Flow panels included in Supplementary Table 1). Cytosolic and intranuclear proteins were detected after permeabilization with the FoxP3 Transcription Factor fixation/permeabilization buffer set (eBioscience Cat. 00-5523-00). Samples were acquired on the FACSymphony instrument (BD Biosciences) using BD FACSDiva software. AccuCheck Counting Beads were used according to manufacturer instructions to calculate absolute cells in samples (Invitrogen; Cat. PCB100). UltraComp™ ebeads (ThermoFisher; Cat. 01-2222-42) and ArC™ Amine Reactive compensation beads were used for compensation. Analysis was performed using FlowJo software version10.10.0.

### IL-15 TRANS-PRESENTATION STAINING

dLN and VT were harvested into RP10 media and minced with scissors prior to a 30 min incubation in DM10 (10% heat-inactivated FBS, 2 mM L-glutamine, 100 U/mL penicillin-streptomycin, 5 mM sodium pyruvate) containing collagenase D (2mg/mL; Sigma Cat.11088858001) and DNAse I (15µg/mL; Sigma Cat.10104159001) at 37 °C with gentle shaking. Tissue samples were then mashed through a 100um strainer and washed with 20mL of RP10 media. VT samples were spun down, and the entire cell pellet was resuspended with fluorescence-activated cell sorting (FACS) buffer into a single-cell suspension, stained, and used for flow cytometric analysis. After staining VT and dLN samples with viability stain (LIVE/DEAD™ Fixable Blue Dead Cell Stain; Cat. L23105) and washing with 1X PBS, samples were stained with L-15 Polyclonal Antibody, Biotin (PeproTech®; Cat. 500-P173BT-25UG) alone at 1:100 in FACS buffer for 30 min at 4C. Samples were washed twice with 200µL of FACS buffer before staining with an extracellular APC panel also containing the secondary antibody, PerCP/Cyanine5.5 Streptavidin (Biolegend; Cat.405214) at 1:500.

### IN VITRO PEPTIDE STIMULATION AND INTRACELLULAR CYTOKINE STAINING

VT and dLN single cell suspensions from experimental and control mice were incubated at 37°C for 4 hours with 10uM HSV-2 gB498-505 peptide (GeneMed Synthesis Inc. Lot# 104865) and 7uM HSV-2 gD290-302 (GeneMed Synthesis Inc. Lot# 106044) diluted in RP10 media with GolgiStop (BD Biosciences Cat. 555029). Unstimulated and PMA/Ionomycin conditions were included as negative and positive controls. After peptide stimulation, cells were washed, pelleted, and stained for flow cytometric analysis of intracellular cytokines using the FoxP3 Transcription Factor fixation/permeabilization buffer set (eBioscience Cat. 00-5523-00) to stain for intracellular IFNγ expression.

### OT-I ADOPTIVE TRANSFERS

CD8+ T cells were isolated from naive Thy1.1 CD45.2 TCR-transgenic OT-I mouse splenocytes using a murine CD8+ T cell isolation kit (Stemcell Tech, Cat. 19853). Isolated cells were washed with PBS and transferred into naive recipient mice via ROI; each mouse received 5e^5^ – 1e^6^ OT-I cells intravenously. 24 hours later, mice were intravaginally infected with 10^5^ PFU of HSV-2 TK-OVA. 7 days post-HSV-2 TK-OVA infection, mice were bled via the submandibular (facial) vein using 5mm Goldenrod lancets. Bleeds were stained using an MHC-I OVA tetramer (conjugated to SA-PE) and congenic markers (Thy1.1, CD45.2) and analyzed using flow cytometry to identify the expanded adoptively transferred population.

### IN VIVO FTY720, TREG DEPLETION, AND MONOCLONAL ANTIBODY DEPLETION

Treg depletion: On days –1 and 0 relative to WT HSV-2 challenge, mice were administered 30µg/kg and 10µg/kg respectively, of diphtheria toxin (DT) via intraperitoneal injection to ablate Treg systemically in FoxP3 DTR mice.

FTY720 treatment: Mice previously infected with either 10^5^ PFU HSV-2 TK-or TK-OVA virus 30 days prior, received 1mg/kg FTY720 (Sigma Cat. SML0700) dissolved in 2% cyclodextrin (ApexBio; Cat.B6413; in sterile PBS) via intraperitoneal (i.p) injection on days –1, 0, 1, and 2 relative to WT HSV-2 challenge. Control mice received vehicle only injections (2% cyclodextrin in PBS).

IL-2 blockade: On day –1 relative to WT HSV-2 challenge, mice were administered 150µg of i.v. anti-IL-2 (Bio X Cell, clone JES6-1A12) or 150µg of i.v. isotype control (Bio X Cell InVivoMAb rat IgG2a isotype control, anti-trinitrophenol). Anti-IL-2 treated mice received 80µg of anti-IL-2 Mab in 10µL intravaginally on day –1, and 40µg intravaginally on days 1 and 2. Control mice received isotype control clones via identical route, concentration, and treatment schedule as experimental mice.

### HISTOLOGY

VTs (including cervix) were carefully harvested into PBS. The tissue was then laid flat in a cassette with a nytex square and submerged in 10% NBF (Sigma; Cat. HT501320) for 3-5 days. Formalin fixed, paraffin embedded samples were then sectioned and H&E stained. Sections were sent to the Fred Hutch Histopathology core to be blindly scored by a pathologist (Amanda Koehne) as previously described^50^. Scores were given to reflect changes in the mucosal epithelium, inflammation within the lamina propria, inflammation within the muscularis, and cellular debris within the vaginal lumen. Inflammatory changes were scored in the mucosal epithelium, lamina propria, and muscularis mucosa, and lumen layers of the VT and then reported as a sum composite. Scoring reflected severity and extent of lesions whereby 0 = no significant change, 1 = minimal, 2 = mild, 3 = moderate, and 4 = severe.

### HSV-2 VIRAL LOAD PCRs

The mouse vaginal canal was carefully swabbed before washing by gently pipetting 50µL of titration buffer in and out. The 50µL of vaginal wash was then added to a screw-top 2mL tube containing 950µL of titration buffer and frozen at-80°C. Vaginal washes were thawed once prior to quantifying HSV-2 viral titers by RT-PCR at UW Virology. DNA was extracted from 200 µl specimen using QIAamp 96 DNA blood kit and eluted into 100 µl of AE buffer (Qiagen).

Real-time Taqman PCR detects HSV gB gene was applied to quantify HSV in the samples^106^. Each 30 µl PCR reaction contains 10 µl of purified DNA, 833 nM primers, 100 nM probe, 15 µl of 2x QuantiTect Multiplex PCR master mix, 0.03 units of UNG. To monitor PCR inhibition, EXO internal control was spiked into all the reactions. The thermocycling conditions are as follows: 50°C for 2 minutes, 95°C for 15 minutes, followed by 45 cycles of 94°C for 1 minute and 60°C for 1 minute. The limit of detection is 3 viral copies/reaction.

### BULK RNA-SEQUENCING

Bulk RNA-sequencing was performed on 100-200 CD4+ FoxP3+ Treg from the dLN and VTs of previously infected and naïve mice. Cells were not pooled between mice, 16 samples were sequenced in total (D28 post-TK-n=4, naïve depo-treated mice n=4). Treg were sorted into directly into SMART-Seq v4 Ultra Low Input lysis buffer (Takara Bio USA) using the BD Symphony S6 and Flash frozen; upon thawing samples underwent reverse transcription followed by PCR amplification to generate full length amplified cDNA. cDNA was then submitted for bulk RNA-seq performed by the Genomics Core at Benaroya Research Institute. Sequencing libraries were constructed using the NexteraXT DNA sample preparation kit with unique dual indexes (Illumina) to generate Illumina-compatible barcoded libraries. Libraries were pooled and quantified using a Qubit® Fluorometer (Life Technologies). Sequencing of pooled libraries was carried out on a NextSeq 2000 sequencer (Illumina) with paired-end 59-base reads (Illumina) with a target depth of 5 million reads per sample. Base calls were processed to FASTQs on BaseSpace (Illumina), and a base call quality-trimming step was applied to remove low-confidence base calls from the ends of reads. The FASTQs were aligned to the GRCm38 mouse reference genome, using STAR v.2.4.2a and gene counts were generated using htseq-count. QC and metrics analysis was performed using the Picard family of tools (v1.134). We received a report including QC, gene counts, and differential gene expression.

**Analysis:** Raw gene counts were analyzed for top differentially expressed genes by the Benaroya Research Institute sequencing core. For comparison of differentially expressed genes, genes with a False Discovery Rate (FDR) of less than 0.1 and an absolute expression fold-change of greater than 1.0 were considered differentially expressed. Volcano plots were visualized using ggplot2 (https://ggplot2.tidyverse.org) and heatmaps were generated in base R (R version 4.5.1).

## STATISTICAL ANALYSES

All statistical analyses were performed using Prism software (GraphPad Software). Statistical significance was determined using unpaired t-tests, one-and two-way ANOVA with ad hoc Tukey’s, Dunn-Šídák, or Dunnett’s multiple comparisons tests, or multiple unpaired t-tests with Welch correction and Holm-Šídák multiple hypothesis testing. For all flow cytometry, virology, and histology studies, each measurement was taken from a distinct sample as biological replicates.

## DATA AVAILABILITY

The sequencing data from this publication have been deposited in the NCBI’s Gene Expression Omnibus and are accessible through the series accession number GEO: GSE318923. *<u>(Editor/Reviewer access token code can be provided upon request)</u>*.

## ACKNOWLEDGEMENTS

We thank all members of the Lund and Prlic labs for their helpful discussion on experimental findings. We also thank Dr. Nicole Potchen, Molly Kanagy, and Anthony Reynolds for their assistance with tissue processing. This work was supported by the following grants from the National Institutes of Health: R01 AI141435 (to J.M.L.), R01 AI172111 (to J.M.L.), and R01 AI123323 (to M.P.). This research was supported by the Benaroya Research Institute Bioinformatics and Genomics Core, Experimental Histopathology Shared Resource, Comparative Medicine Shared Resource, and Immune Monitoring Shared Resource, RRID: SCR_022612, of the Fred Hutch/University of Washington/Seattle Children’s Cancer Consortium (P30 CA015704). The Experimental Histopathology Shared Resource equipment is supported by a grant from the M.J. Murdock Charitable Trust grant SR-202221337. ICT was supported by the Seattle Chapter ARCS Foundation Fellowship and T32 AI007140, and LW was supported by T32 AI083203.

## AUTHOR CONTRIBUTIONS

ICT, JBG, JLS, BRT, LW, and TA conducted experiments. ICT, JBG, JLS, BRT, MQP, and ALK analyzed data. KRJ, MP, and JML provided supervision. ICT and JML designed the research study and wrote the first draft of the manuscript. All authors edited and approved the manuscript.

## Supplemental Figure Legends

**Supplementary Figure 1 (S1). Treg phenotyping in the VT and dLN after primary HSV-2 infection.** Female FoxP3^eGFP^ mice were intravaginally infected with 10^5^ PFU of thymidine kinase-deficient HSV-2 5-7 days after subcutaneous (neck ruff) administration of Depo Provera (2mg/kg in sterile PBS). VT and dLN were collected at 7 and 90 days post-infection and processed for flow cytometric analysis, with naïve depo-treated mice included as day 0 uninfected controls. (**A**) Frequencies of Treg expression for different markers of activation and immunosuppressive function in the VT (top) and dLN (bottom) at day 0, 7, and 90. Data shown are combined from two experiments per timepoint (n= 3-8 mice per group). Error bars represent the mean and SD. Statistical significance was determined by One-Way ANOVA and Dunnett’s multiple comparisons test. (**B**) Representative staining for Treg expression of each marker of activation and immunosuppressive function at days 0, 7, and 90.

**Supplementary Figure 2 (S2). T cell phenotypic marker frequencies and counts upon secondary infection.** Depo Provera-treated Female FoxP3^WT^ and FoxP3^DTR^ mice were vaginally infected with 10^5^ PFU of attenuated HSV-2 TK-and allowed to recover for 28-30 days. Mice were again treated with Depo Provera 5-7 days before WT HSV-2 (10^4^ PFU) challenge and treated intraperitoneally with diphtheria toxin (DT) to ablate Treg prior to infection (FoxP3^DTR^ mice, Treg-depleted group, Treg-). A dose of 30 μg/kg was administered the day before WT infection, followed by a dose of 10 μg/kg on d0. VT and dLN were harvested at 3 days p.c. and characterized via flow cytometry. (**A**) Absolute number of vaginal and dLN FoxP3+ CD4+ Treg. (**B**) Absolute number of CD8+T cells in Treg-depleted and sufficient mice after WT HSV-2 challenge. (**C**) Absolute number of VT Ki67+ CD44+ CD8 T cells. (**D**) Frequency (left) and absolute number of dLN Ki67+ CD44+ CD8 T cells in Treg-depleted and sufficient mice after WT HSV-2 challenge. (**E**) Absolute number of VT GrzmB+ CD44+ CD8 T cells. (**F**) Frequency (left) and absolute number (right) of dLN GrzmB+ CD44+ CD8 T cells. (**G**) Representative staining for CD45.2 I.V. label and CD69 vs. CD103 gating for Trm. (**H**) Absolute number of VT Ki67+ CD44+ CD8 Trm (defined as CD69+ CD103+ I.V.-label negative). (**I**) Absolute number of VT GrzmB+ CD44+ CD8 Trm. (**J**) Female FoxP3^WT^ and FoxP3^DTR^ mice were infected and treated with DT as in Figure 3. In addition, mice were treated with 1mg/kg FTY720 via intraperitoneal injections on days-1, 0, and 2 with respect to WT HSV-2 challenge. Control mice were treated with vehicle (2% cyclodextrin in sterile PBS).

Absolute number of Ki67+ CD44+ I.V.-label negative CD8 T cells in FTY720 and vehicle control treated HSV-challenged mice. (**K**) Absolute number of VT Ki67+ CD44+ CD8 Trm in FTY720 and vehicle-treated control HSV-challenged mice. Representative data from two experiments (n= 4-5 mice per group) are shown. Statistical significance was determined by One-Way ANOVA and Dunn-Šídák multiple comparisons tests in A-F and H-K.

**Supplementary Figure 3 (S3). HSV-2 gB-speciQc CD8 T cells.** Female FoxP3^WT^ and FoxP3^DTR^ mice were infected and treated with DT as in Figure 3. (**A**) Frequency (left) and absolute number (right) of gB tetramer+ CD44+ CD8 T cells in dLN of WT HSV-2-challenged Treg-depleted and sufficient mice. (**B**) Representative staining of dLN gB tetramer+ CD44+ CD8 T cells. (**C**) Frequency (left) and absolute number (right) of dLN Ki67+ CD44+ gB tetramer+ CD8 T cells. (**D**) Frequency (left) and number (right) of VT Ki67+ CD44+ gB tetramer+ CD8 T cells. (**E**) Frequency of VT CD44+ gB tetramer+ Trm. (**F**) Frequency (left) and absolute number (right) of VT Ki67+ CD44+ gB tetramer+ Trm. (**G**) Female FoxP3^WT^ and FoxP3^DTR^ mice were infected and treated with DT as in Figure 3. In addition, mice were treated with 1mg/kg FTY720 via intraperitoneal injections on days-1, 0, and 2 with respect to WT HSV-2 challenge. Control mice were treated with vehicle (2% cyclodextrin in sterile PBS).

Frequency and absolute number of VT Ki67+ CD44+ gB tetramer+ I.V.-label negative CD8 T cells in FTY720 and vehicle control treated HSV-challenged mice. (**H**) Frequency of dLN GrzmB+ CD44+ gB tetramer+ CD8 T cells. (**I**) Frequency (left) and absolute number (right) of dLN GrzmB+ NKG2D+ gB tetramer-negative CD44+ CD8 T cells. Data combined from two experiments for each plot (n= 4-5 mice per group). Error bars represent the mean and SD. Statistical significance was determined by One-Way ANOVA and Dunn-Šídák multiple comparisons tests in A, C-I.

**Supplementary Figure 4 (S4). Cytotoxic bystander-activated OT-I cells and *in vivo* IL-2 Mab neutralization.** 500,000-1 million OT-Is were adoptively transferred into female FoxP3^WT^ and FoxP3^DTR^ mice 24 hours before vaginal infection with 10^5^ PFU of HSV-2 TK-OVA and allowed to recover for 28-30 days. Mice were again treated with Depo Provera 5-7 days before WT HSV-2 (10^4^ PFU) challenge and treated intraperitoneally with diphtheria toxin (DT) to ablate Treg prior to infection (FoxP3^DTR^ mice, Treg-depleted group, Treg-). VT and dLN were collected d3 p.c. and assessed via flow cytometry. (**A**) Absolute number of dLN Thy1.1+ CD44+ OT-1 CD8 T cells. (**B**) Frequency (left) and absolute number (right) of bystander-activated dLN GrzmB+ NKG2D+ Thy1.1+ CD44+ OT-I CD8 T cells. (**C**) Frequency (left) and absolute number (right) of VT Ki67+ Thy1.1+ CD44+ OT-I CD8 T cells. (**D**) Mice were administered both intravenous (150µg) and intravaginal (80µg) anti-IL-2 monoclonal antibody (clone JES6-1A12) or rat IgG2a isotype control on day-1 with respect to WT challenge. 40µg of anti-IL-2 or isotype control was administered intravaginally d1 and 2. VT and dLN were collected 3 days p.c. and assessed via flow cytometry. Shown in plots are the frequency (left) and absolute number (right) of total dLN GrzmB+ CD44+ CD8 T cells from WT HSV-2 challenged Treg-depleted or sufficient mice after treatment with anti-IL-2 or isotype control.

(**E**) Absolute number of VT (left) and dLN (right) gB tetramer+ CD44+ CD8 T cells from WT HSV-2 challenged Treg-depleted or sufficient mice after treatment with anti-IL-2 or isotype control. (**F**) Frequency (left) and absolute number (right) of dLN GrzmB+ gB tetramer+ CD44+ CD8 T cells from WT HSV-2-challenged Treg-depleted or sufficient mice after treatment with anti-IL-2 or isotype control. (**G**) Absolute number of VT Ki67+ CD44+ gB tetramer+ CD8 T cells from WT HSV-2 challenged Treg-depleted or sufficient mice after treatment with anti-IL-2 or isotype control. (**H**) Frequency and absolute number of bystander-activated dLN NKG2D+ GrzmB+ CD44+ OT-I CD8 T cells from WT HSV-2-challenged Treg-depleted or sufficient mice after treatment with anti-IL-2 or isotype control. Data shown are combined from two experiments (n=3-5 mice per group). Samples in which the previous gate contained fewer than 25 events were excluded from analysis. Error bars represent the mean and SD. Statistical significance was determined by One-Way ANOVA and Dunn-Šídák multiple comparisons tests.

**Supplementary Figure 5 (S5). IL-15 trans-presentation by APC.** Female FoxP3^WT^ and FoxP3^DTR^ mice were infected and treated with DT as in Figure 3. VT and dLN were collected d3 p.c. and assessed via flow cytometry. (**A**) Shown is the gating strategy used to identify APC in VT and dLN tissues (representative example from VT sample). (**B**) Absolute number of of IL-15 trans-presenting myeloid cells (Ly6G-CD19+ CD3e-CD45+) in VT from WT HSV-2-challenged Treg-depleted or sufficient mice. (**C**) Absolute number of IL-15 trans-presenting myeloid cells (Ly6G-CD19+ CD3e-CD45+) in dLN from WT HSV-2 challenged Treg-depleted or sufficient mice. Data shown in B and C are representative of two experiments with 4-5 mice per group. (**D**) Mice were administered both intravenous (150µg) and intravaginal (80µg) anti-IL-2 monoclonal antibody (clone JES6-1A12) or rat IgG2a isotype control on day-1 with respect to WT challenge. 40µg of anti-IL-2 or isotype control was administered intravaginally d1 and 2. VT and dLN were collected 3 days p.c. and assessed via flow cytometry. Shown is the absolute numbers of vaginal IL-15+ monocytes, (**E**) IL-15+ macrophages, and (**F**) IL-15+ dendritic cells from WT HSV-2-challenged Treg-depleted or sufficient mice after treatment with anti-IL-2 or isotype control. (**G**) Frequency of dLN IL-15+ monocytes, (**H**) IL-15+ macrophages, and (**I**) IL-15+ dendritic cells from WT HSV-2-challenged Treg-depleted or sufficient mice after treatment with anti-IL-2 or isotype control. (**J**) Absolute number of IL-15+ dLN monocytes, (**K**) IL-15+ macrophages, (**L**) IL-15+ dendritic cells from WT HSV-2-challenged Treg-depleted or sufficient mice after treatment with anti-IL-2 or isotype control. Error bars represent the mean and SD. Statistical significance was determined by One-Way ANOVA and Dunn-Šídák multiple comparisons tests for B-L. **(M**) Representative staining of the IL-2R components in VT APC (CD25, CD122, CD132) about 33 days after primary vaginal infection with TK-HSV-2.

---

**Supplementary Table 1.**
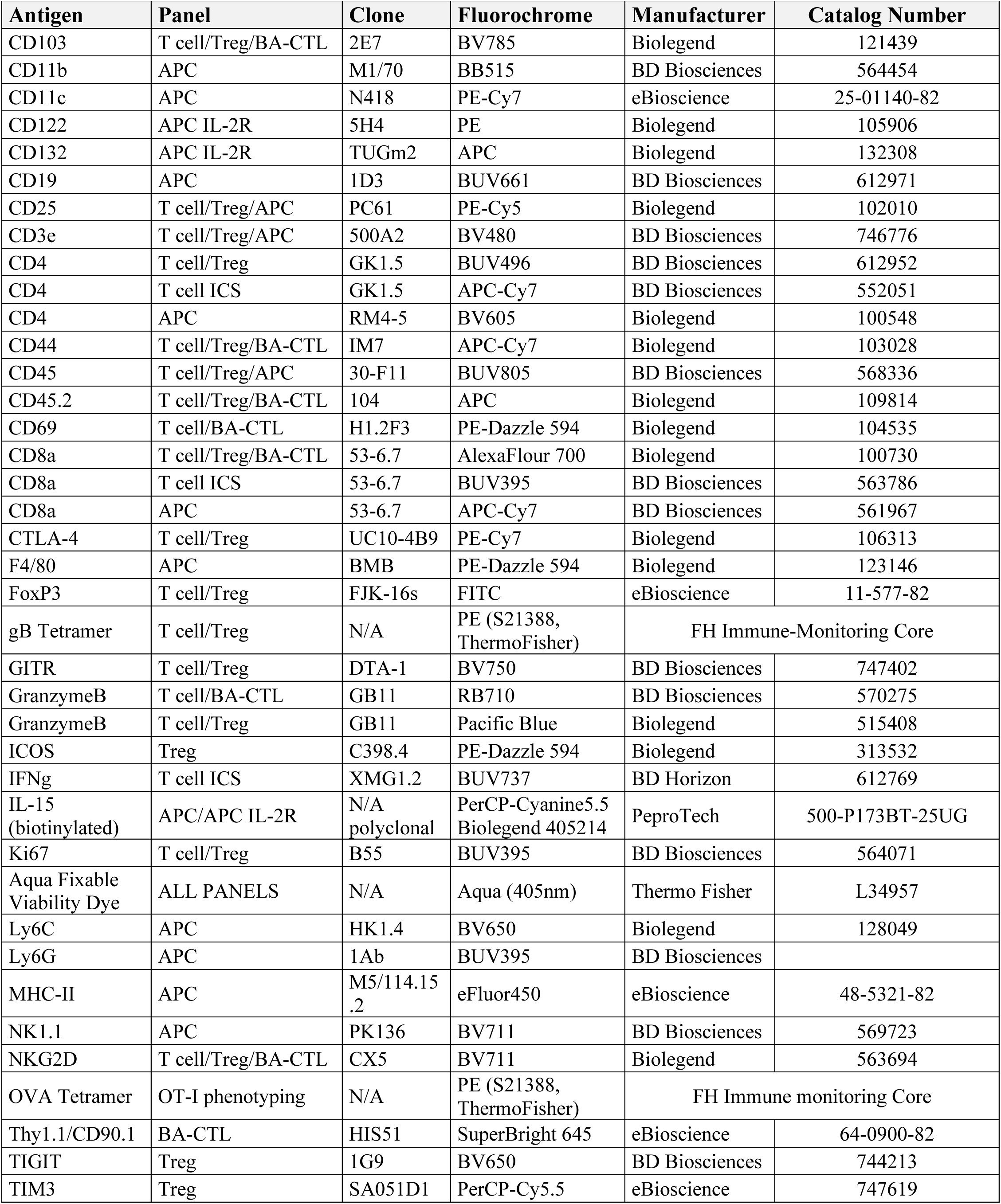
Flow Cytometry Antibody Details.

| **Antigen** | **Panel** | **Clone** | **Fluorochrome** | **Manufacturer** | **Catalog Number** |
| --- | --- | --- | --- | --- | --- |
| CD103 | T cell/Treg/BA-CTL | 2E7 | BV785 | Biolegend | 121439 |
| CD11b | APC | M1/70 | BB515 | BD Biosciences | 564454 |
| CD11c | APC | N418 | PE-Cy7 | eBioscience | 25-01140-82 |
| CD122 | APC IL-2R | 5H4 | PE | Biolegend | 105906 |
| CD132 | APC IL-2R | TUGm2 | APC | Biolegend | 132308 |
| CD19 | APC | 1D3 | BUV661 | BD Biosciences | 612971 |
| CD25 | T cell/Treg/APC | PC61 | PE-Cy5 | Biolegend | 102010 |
| CD3e | T cell/Treg/APC | 500A2 | BV480 | BD Biosciences | 746776 |
| CD4 | T cell/Treg | GK1.5 | BUV496 | BD Biosciences | 612952 |
| CD4 | T cell ICS | GK1.5 | APC-Cy7 | BD Biosciences | 552051 |
| CD4 | APC | RM4-5 | BV605 | Biolegend | 100548 |
| CD44 | T cell/Treg/BA-CTL | IM7 | APC-Cy7 | Biolegend | 103028 |
| CD45 | T cell/Treg/APC | 30-F11 | BUV805 | BD Biosciences | 568336 |
| CD45.2 | T cell/Treg/BA-CTL | 104 | APC | Biolegend | 109814 |
| CD69 | T cell/BA-CTL | H1.2F3 | PE-Dazzle 594 | Biolegend | 104535 |
| CD8a | T cell/Treg/BA-CTL | 53-6.7 | AlexaFlour 700 | Biolegend | 100730 |
| CD8a | T cell ICS | 53-6.7 | BUV395 | BD Biosciences | 563786 |
| CD8a | APC | 53-6.7 | APC-Cy7 | BD Biosciences | 561967 |
| CTLA-4 | T cell/Treg | UC10-4B9 | PE-Cy7 | Biolegend | 106313 |
| F4/80 | APC | BMB | PE-Dazzle 594 | Biolegend | 123146 |
| FoxP3 | T cell/Treg | FJK-16s | FITC | eBioscience | 11-577-82 |
| gB Tetramer | T cell/Treg | N/A | PE (S21388, ThermoFisher) | FH Immune-Monitoring Core |  |
| GITR | T cell/Treg | DTA-1 | BV750 | BD Biosciences | 747402 |
| GranzymeB | T cell/BA-CTL | GB11 | RB710 | BD Biosciences | 570275 |
| GranzymeB | T cell/Treg | GB11 | Pacific Blue | Biolegend | 515408 |
| ICOS | Treg | C398.4 | PE-Dazzle 594 | Biolegend | 313532 |
| IFNg | T cell ICS | XMG1.2 | BUV737 | BD Horizon | 612769 |
| IL-15 (biotinylated) | APC/APC IL-2R | N/A polyclonal | PerCP-Cyanine5.5 Biolegend 405214 | PeproTech | 500-P173BT-25UG |
| Ki67 | T cell/Treg | B55 | BUV395 | BD Biosciences | 564071 |
| Aqua Fixable Viability Dye | ALL PANELS | N/A | Aqua (405nm) | Thermo Fisher | L34957 |
| Ly6C | APC | HK1.4 | BV650 | Biolegend | 128049 |
| Ly6G | APC | 1Ab | BUV395 | BD Biosciences |  |
| MHC-II | APC | M5/114.15.2 | eFluor450 | eBioscience | 48-5321-82 |
| NK1.1 | APC | PK136 | BV711 | BD Biosciences | 569723 |
| NKG2D | T cell/Treg/BA-CTL | CX5 | BV711 | Biolegend | 563694 |
| OVA Tetramer | OT-I phenotyping | N/A | PE (S21388, ThermoFisher) | FH Immune monitoring Core |  |
| Thy1.1/CD90.1 | BA-CTL | HIS51 | SuperBright 645 | eBioscience | 64-0900-82 |
| TIGIT | Treg | 1G9 | BV650 | BD Biosciences | 744213 |
| TIM3 | Treg | SA051D1 | PerCP-Cy5.5 | eBioscience | 747619 |

