## Supplemental Figures 1-5 for "Regulatory T cells restrain cytotoxic and bystander CD8 T cells without compromising antigen-driven memory in mucosal tissue"

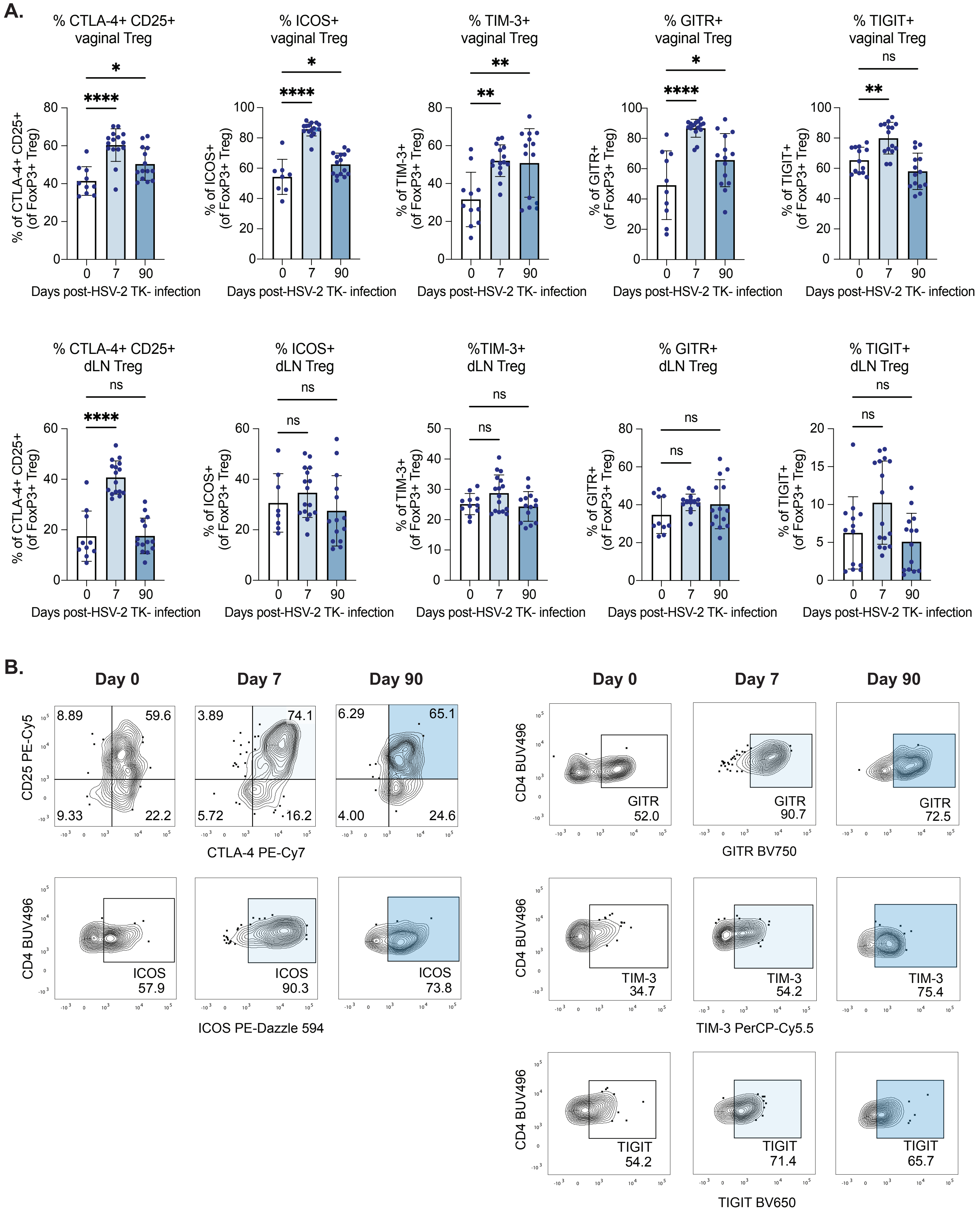


***Supplementary Fig. 1***

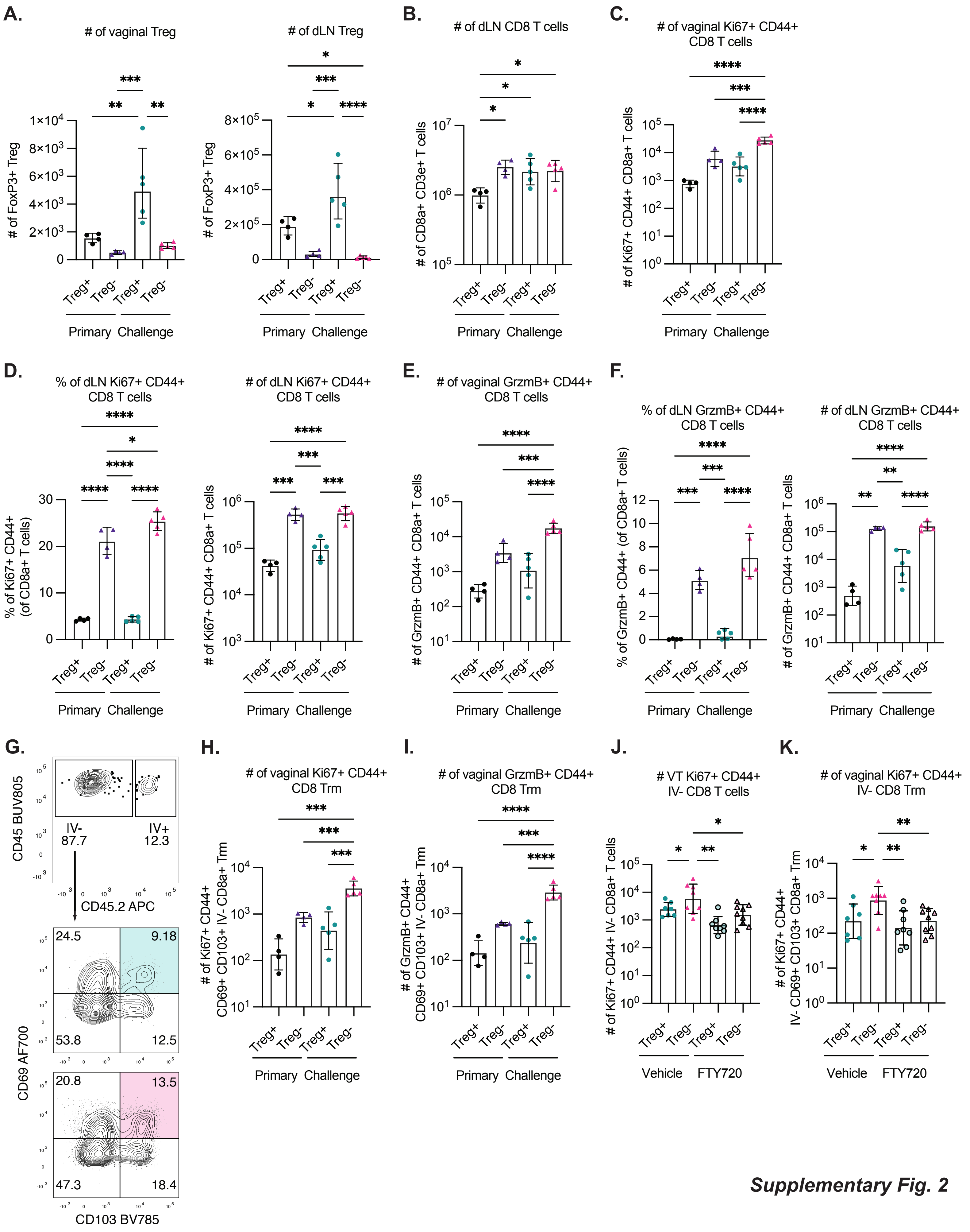


**Supplementary Fig. 2**

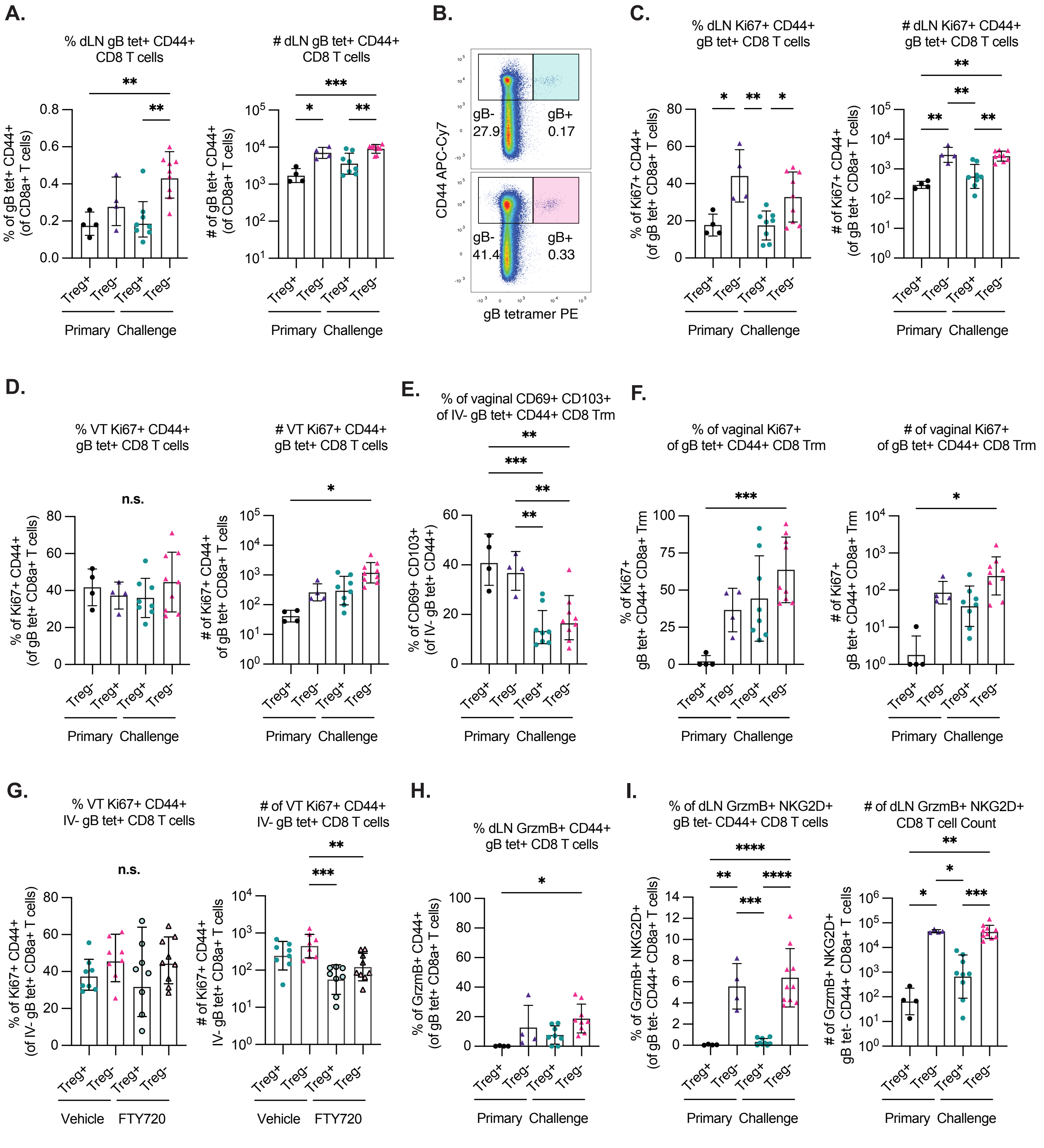


*Supplementary Fig. 3*

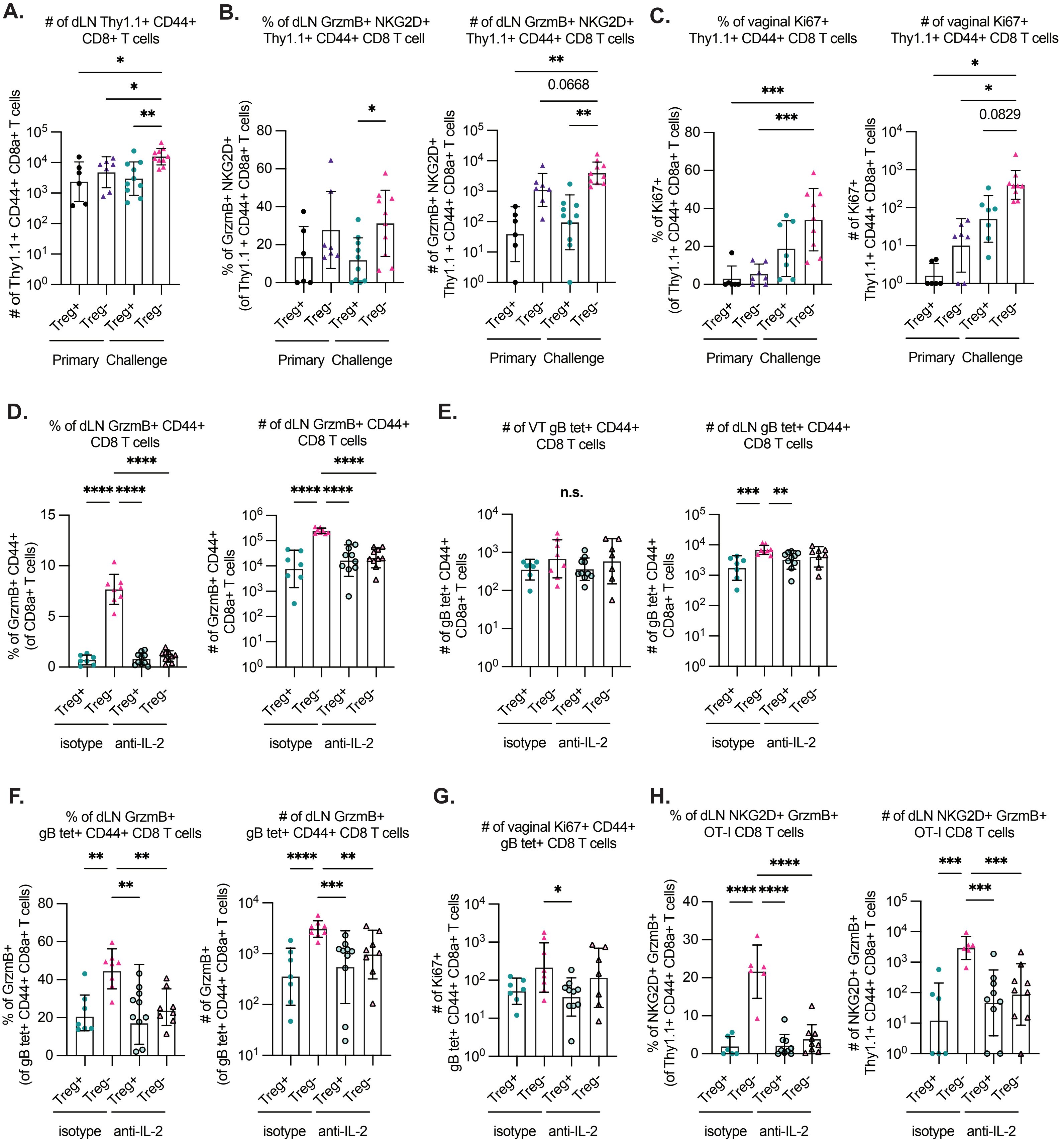


***Supplementary Fig. 4***

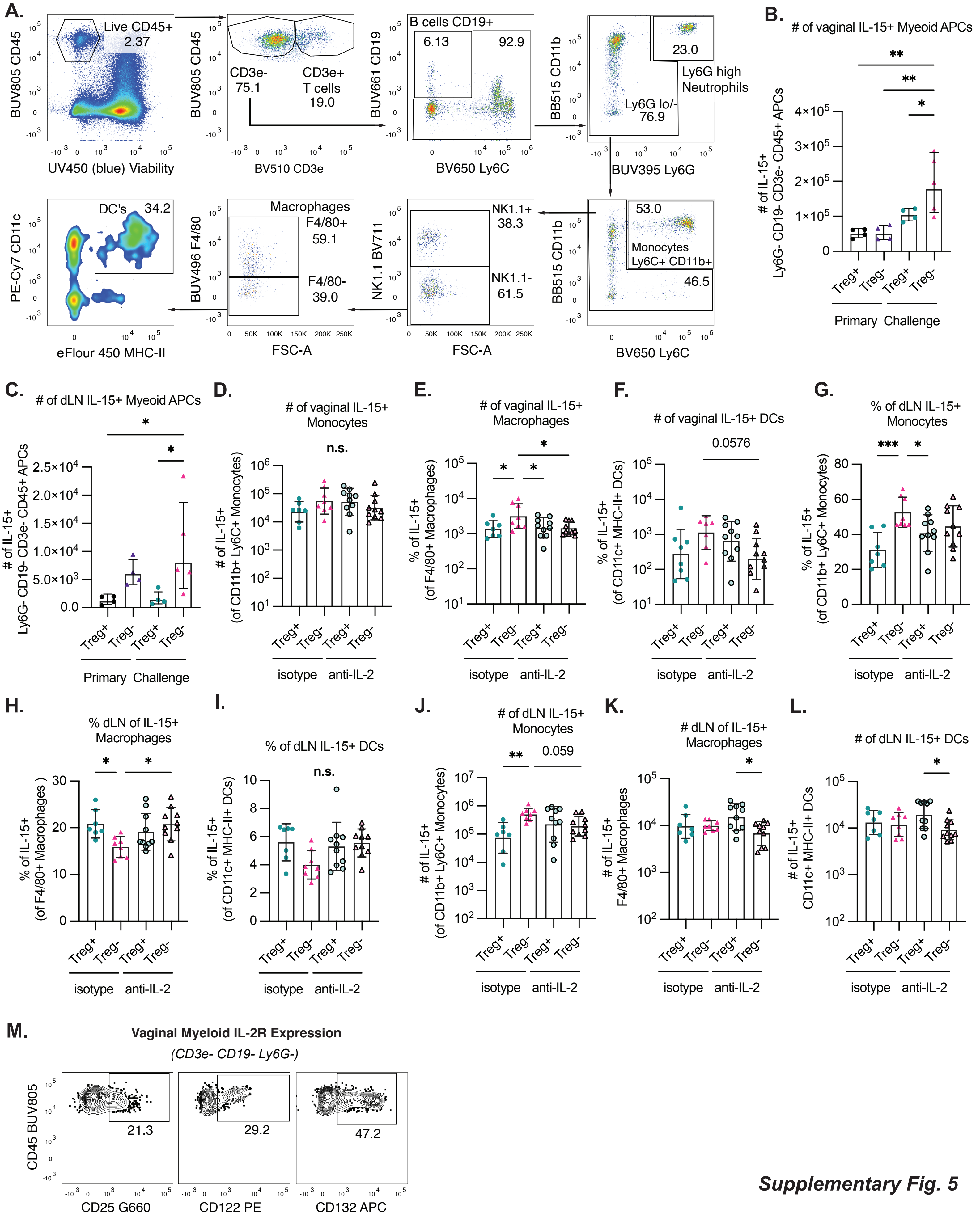


***Supplementary Fig. 5***
